# Enhancer lncRNA LOC730338 modulates BCR signaling and immune evasion in lymphoma by regulating RNA homeostasis

**DOI:** 10.1101/2024.12.10.627805

**Authors:** Luciano Cascione, Francesca Guidetti, Sunandini Ramnarayanan, Andrea Rinaldi, Filippo Spriano, Alex Zadro, Chiara Tarantelli, Serena Zambarbieri, Nicolas Munz, Simone Avesani, Alberto J. Arribas, Rosalba Giugno, Rory Johnson, Francesco Bertoni, Sara Napoli

## Abstract

Chronic antigenic stimulation is a central factor in the development of marginal zone lymphoma (MZL). While the pharmacological inhibition of the B-cell receptor (BCR) signaling by Bruton’s tyrosine kinase (BTK) inhibitors is initially effective, the development of resistance remains a challenge in treating MZL and other B-cell malignancies. Enhancer activation remodeling is a key epigenetic mechanism that enables tumor adaptation during therapy. The most active regulatory regions cluster in super-enhancers and produce enhancer RNAs (eRNAs), a class of unstable noncoding transcripts that primarily serve as scaffolds for chromatin looping. However, when stabilized, these eRNAs can evolve into long noncoding RNA (lncRNAs) with distinct functions.

To investigate enhancer-associated long non-coding RNAs (elncRNAs) involved in shaping BCR pathway dependence, we conducted a CRISPR interference (CRISPRi) screen in MZL cells. We identified LOC730338, an elncRNA linked to A-to-I RNA editing, which we renamed ADARreg. ADARreg renders tumor cells refractory to BCR pathway inhibition by modulating ADAR2 nuclear translocation and altering RNA modification patterns in key regulatory isoforms through coordinated control of RNA stability and localization, as revealed through subcellular direct RNA sequencing (DRS). In addition, ADARreg induces an immune-suppressive transcriptional program, increasing the production of inhibitory cytokines and receptors that diminish NK cell-mediated cytotoxicity.

Together, these findings uncover a novel role for elncRNAs in orchestrating immune evasion and provide a potential therapeutic strategy to overcome resistance in lymphoma and other immune-related diseases.

**Graphical abstract:** 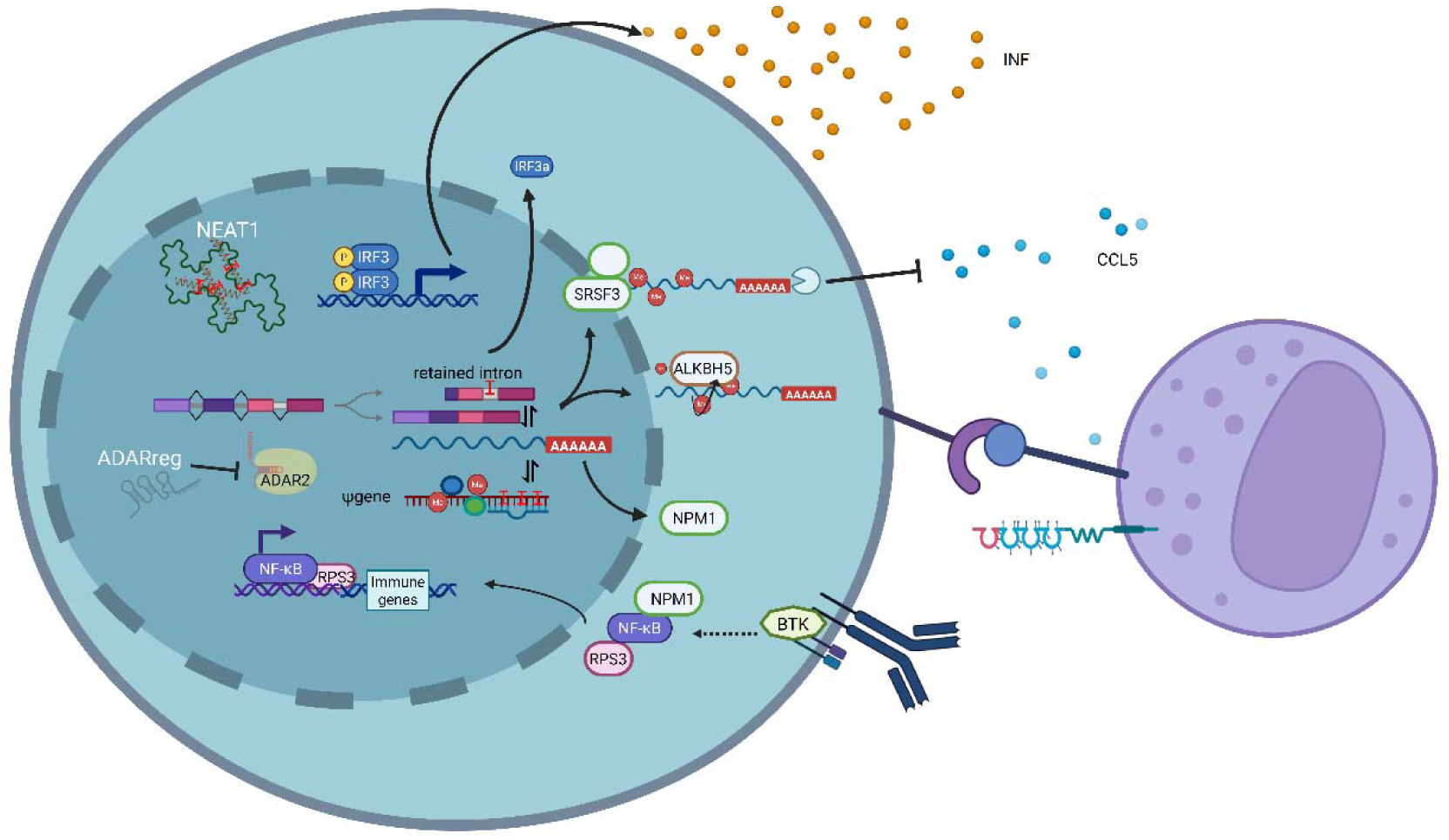

**Highlights:**

- The enhancer-associated long non-coding RNA ADARreg modulates RNA editing and alters the stability and localization of key immune-related transcripts, thereby enabling tumor immune evasion.
- ADARreg has prognostic and therapeutic relevance, linking resistance mechanisms and immune modulation, which supports the development of RNA-based therapeutic strategies in cancer.

## INTRODUCTION

Marginal zone Lymphoma (MZL) derives from MZ B cells under chronic inflammation and prolonged B cell receptor (BCR) activation from persistent infections or autoimmune diseases ^1^. A hallmark of MZL is the interaction of tumor cells with the microenvironment, a complex network of various cell types, including T cells, which communicate through soluble factors like cytokines and chemokines ^2^. BCR signaling plays a crucial role in shaping these interactions and activating downstream pathways like NF-κB to sustain lymphoma growth ^3,4^. Indeed, the pharmacological inhibition of BCR signaling, particularly through Bruton’s tyrosine kinase (BTK) inhibition with the first-in-class ibrutinib and newer agents, is an effective therapeutic option for MZL patients as well as for other B-cell neoplasms and autoimmune disorders ^5,6^. However, resistance frequently develops, sustained by epigenetic reprogramming ^7–11^. The specific regulators driving this transcriptional plasticity remain incompletely understood, with growing attention focusing on non-coding RNAs (ncRNAs), particularly enhancer-associated long non-coding RNAs (elncRNAs), which play key roles in regulating gene expression ^12,13^. ElncRNAs are transcribed from enhancer regions and function as tissue-specific transcriptional regulators ^14^. They derive from super-enhancers (SEs), highly acetylated enhancer clusters that control lineage-defining gene programs ^13,15^. Increasing evidence implicates elncRNAs in cancer ^16^, where their enhancer-derived origin places them at key nodes of oncogenic transcriptional networks. By stabilizing protein complexes *in cis* or acting *in trans* to influence nuclear and cytoplasmic processes, elncRNAs can promote tumor progression and therapeutic resistance ^17,18^. Their restricted expression patterns, tight linkage to specific enhancer landscapes, and capacity to modulate key signaling pathways collectively make elncRNAs attractive candidates for mechanistic investigation and potential therapeutic intervention ^19^.

Here, we aimed to identify elncRNAs modulating the transcriptional reprogramming in ibrutinib-resistant splenic MZL (SMZL) via a screen based on the Clustered Regularly Interspaced Short Palindromic Repeats Interference (CRISPRi) technology. Among the candidates, the lncRNA LOC730338 (AC011294.3/ENSG00000233539) emerged due to its previously identified role in regulating A-to-I RNA editing in B cells ^20^. We show that this transcript likely functions as a negative regulator of ADAR2 activity. Based on this, we renamed it ADARreg and demonstrated its role in modulating the BCR pathway and its immune-suppressive capacity. Using direct RNA sequencing (DRS) of subcellular fractions, we uncovered a complex regulation of RNA homeostasis, including RNA stability and trafficking, orchestrated by ADARreg.

## RESULTS

### Acquired ibrutinib-resistance in SMZL cells is associated with changes in super-enhancer activation

To investigate the mechanism by which MZL cells become BCR-independent and stop responding to anti-BCR pathway treatment, we utilized an *in vitro* model previously developed in our laboratory ^21^. The VL51-Ibru is a derivative of the VL51, a *bona fide* SMZL cell line ^22^, in which a nine-month exposure to increasing concentrations of the BTK inhibitor ibrutinib has led to secondary resistance to BTK inhibitors and other BCR pathway-targeting drugs, such as the PI3K inhibitors idelalisib, duvelisib, and copanlisib ^21^. *In vitro* BCR stimulation revealed that, in the ibrutinib-resistant VL51-Ibru, NF-κB signaling exhibited reduced BCR dependency compared to parental cells (Fig. S1A).

Since VL51-Ibru has no DNA mutations in BCR pathway genes that are recurrently mutated in patients (including *BTK* and *PLCG2*), as reported in 21, we studied the epigenetic remodeling of the resistant and parental cells using ChIP-sequencing to profile histone modifications (H3K27ac, H3K4me1, and H3K4me3). We identified 33,264 H3K27ac peaks: 20,950 common, 2,830 unique to VL51, and 541 unique to VL51-Ibru (Fig. 1A, Table S1). Enhancers were defined by the concomitant presence of H3K27ac and H3K4me1 and the absence of H3K4me3. Compared to its parental counterpart, VL51-Ibru lost 1,329 enhancers and gained 144 new ones (Fig. 1A, B). Utilizing RNA-seq data from parental and derivative cells, we examined the genes located within 40 kb of these differentially activated enhancers. Loss of enhancers was linked to the downregulation of key genes, including FOXO1, CDKN2A, CDKN2B, and MAP3K14, with the most notable decrease observed in NLRP4, a pattern-recognition receptor (PRR) involved in immune activation (Fig.1C) ^23^. A few genes near newly activated enhancers were significantly upregulated in resistant cells (Fig.1C). These findings suggest that, in addition to regulating target genes, newly activated enhancers may function through the non-coding transcripts associated with them.

**Figure 1.**
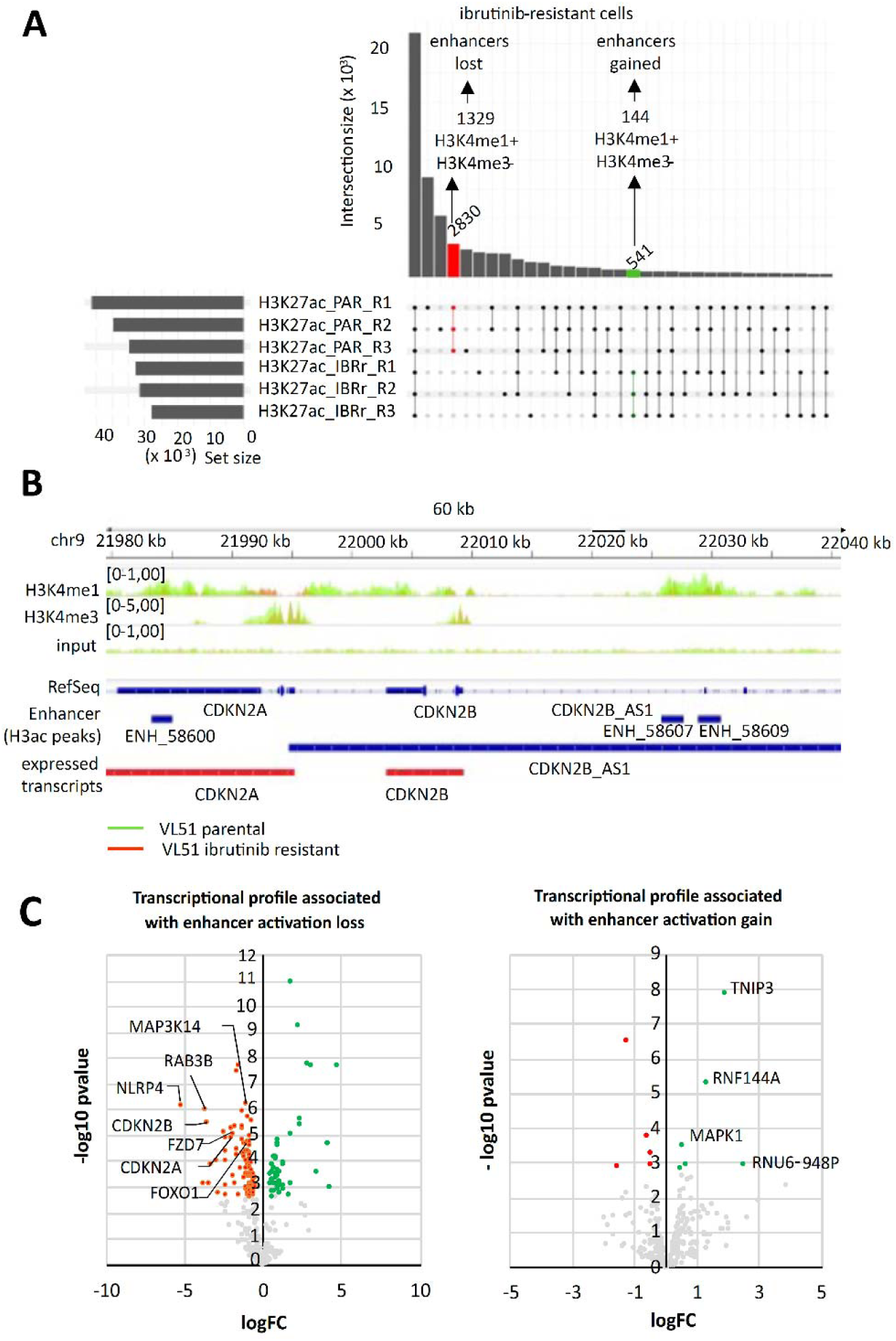
Ibrutinib-resistance is associated with changes in super-enhancer activation. (A) ChIP-seq H3k27ac peak counts in biological replicates of parental VL51 and VL51-Ibru (IBRr). Total peaks (horizontal bars) and robust differentially enriched peaks in VL51 and VL51-Ibru are shown (vertical bars, red and green, respectively). The indicated enhancer numbers consider H3K4me1 peak presence and H3K4me3 absence. (B) Example of an enhancer lost in VL51-Ibru near CDKN2A/B. Tracks show H3K4me1, H3K4me3, and input from VL51 (green) and VL51-Ibru (red), with H3K27ac peaks and differentially expressed transcripts highlighted. (C) Volcano plots of genes within 40 kb of enhancers lost (left) or gained (right) in VL51-Ibru cells. Orange and green denote significantly down- and upregulated genes; selected genes are labeled.

### Identification of elncRNAs associated with ibrutinib resistance

Super-enhancers (SEs) are densely bound by transcription factors and highly acetylated. They control the correct temporal and tissue-specific gene expression and are the primary source of elncRNAs^14,24^. Using the ROSE algorithm ^25^, we identified 1,345 SEs from H3K27Ac ChIP-Seq data. (Fig.S1B). To determine how many SEs were associated with elncRNA production, we reconstructed the transcriptomes of VL51 and VL51-Ibru cells using total RNA-Seq profiling. We included VL51-Ide, a VL51 derivative resistant to PI3Kδ inhibitors ^11^, to expand our analysis beyond BTK inhibition resistance. By analyzing ribosomal RNA-depleted libraries, we detected polyadenylated and non-polyadenylated transcripts and identified novel and known transcripts associated with SEs (Fig. S1B). We defined elncRNAs as any transcript detectable within an active SE in either parental or resistant cells. We assigned a transcription start site (TSS) based on the FANTOM project CAGE dataset ^26^ (Fig. S1B). We excluded transcripts annotated as protein-coding genes but included antisense or sense intronic transcripts. We identified 427 elncRNAs, including 74 novel transcripts not listed in GENCODE (Fig.S1B). Of these, 13% (45 elncRNAs) were lost, and 9% (30 elncRNAs) were gained in the VL51-Ibru cells (Fig.S1B, Table S2).

### Functional screening of elncRNAs linked to BCR blockade

To identify elncRNAs that mediate or oppose BCR blockade, we performed a genome-wide CRISPRi screen in both parental and ibrutinib-resistant cells (Fig.2A). We designed a custom library of paired guide RNAs (pgRNAs) that targeted 255 annotated and 57 novel elncRNAs (Fig.2B). The library also included pgRNAs targeting 225 long intergenic ncRNAs (lincRNAs) differentially expressed between the two cell lines, 68 essential genes as positive controls, and 33 non-expressed transcripts as negative controls. Altogether, the library covered 660 transcripts (Fig.2B, Table S2). The pgRNAs were designed to target regions within -300 to +300 base pairs around the TSSs by a dead Cas9 fused with the repressor ZIM3 (dCas9-KRAB-ZIM3) (Fig.2A, S1C), to transcriptionally repress avoiding DNA damage ^27^.

**Figure 2.**
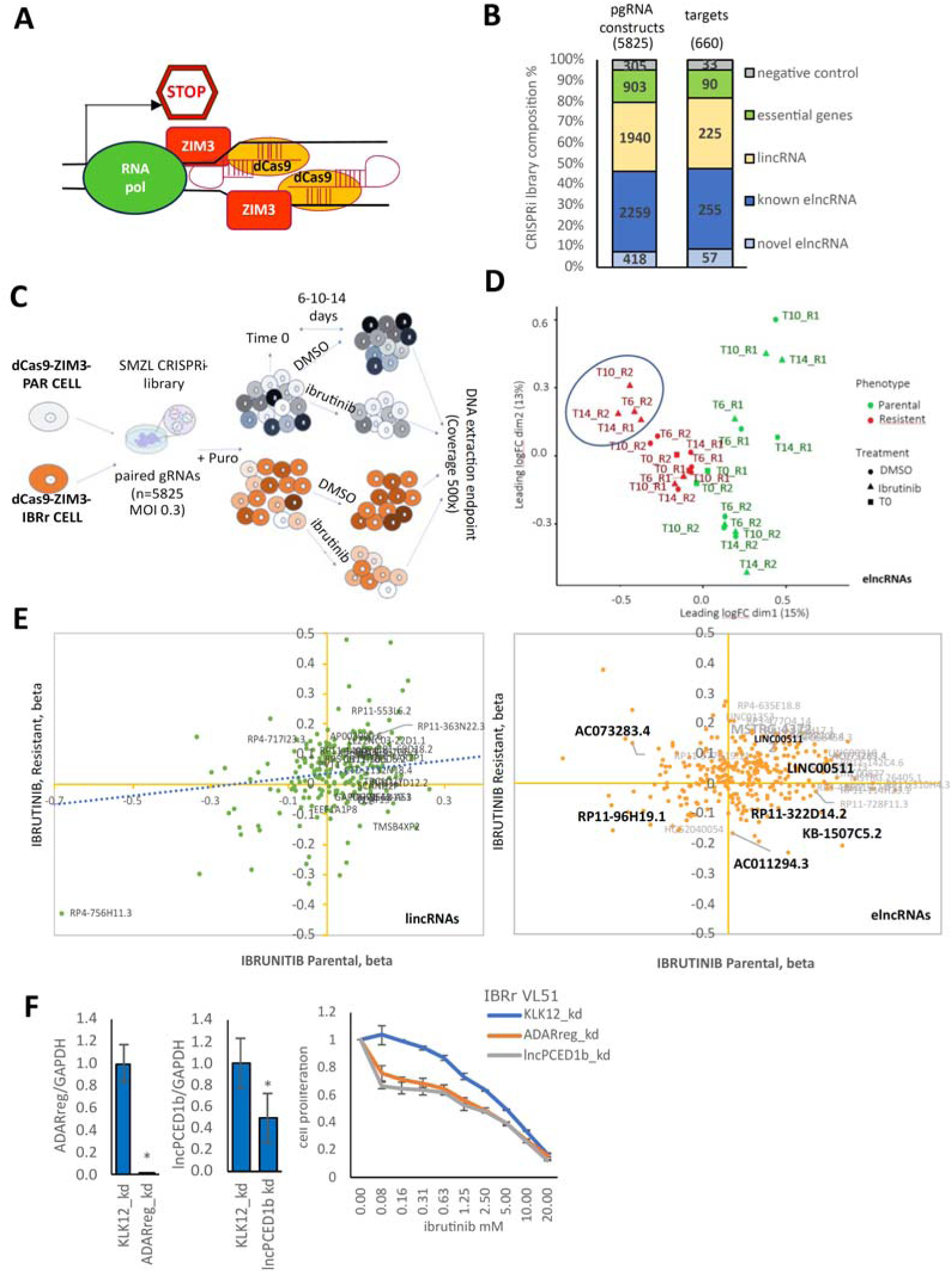
Functional screening of elncRNAs linked to BCR blockade. (A) CRISPRi strategy scheme. (B) Composition of the CRISPRi library. (C) Workflow of the CRISPRi screen. (D) MDS plots of samples at baseline (T0) or after 6, 10, and 14 days. VL51 (green) and VL51-Ibru (red) exposed to ibrutinib (triangles) or vehicle (circles) are shown. VL51-Ibru treated with ibrutinib cluster together. (E) Scatter plots of β-values reflecting dropout of lincRNAs (left) or elncRNAs (right) in VL51 versus VL51-Ibru after 14 days of ibrutinib. Statistically significant hits (FDR < 0.2) are labeled; selected candidates are highlighted. (F) Expression of ADARreg and lncPCED1b, and ibrutinib sensitivity of monoclonal knockdown lines compared with KLK12 controls (n=3,* P <0.05).

To evaluate the impact of elncRNA silencing on BCR blockade, after elncRNA CRISPRi library infection (Fig.S1D), parental and resistant cells were treated with either ibrutinib or DMSO for two weeks (Fig.2C). We analyzed two replicates per condition at three time points (6, 10, and 14 days) compared to time 0 (Fig.2C). The effects of specific transcripts on cell survival and BCR dependency, assessed by ibrutinib sensitivity, were measured through pgRNAs enrichment overtime, represented by β score (Fig.2D, Table S3). Positive β values indicate that transcripts impaired proliferation or increased drug sensitivity, negative β values indicate the opposite. The high β correlation for positive controls across replicates validated the robustness of our approach (Fig.S1E).

The β correlation for elncRNAs targets was consistent, albeit higher in parental cells, possibly reflecting greater heterogeneity in the resistant population (Fig.S1F). Multi-dimensional scaling (MDS) plots showed that lincRNA silencing differentiated the two cell lines, especially after 14 days (Fig. S1G). Similarly, elncRNA silencing discriminated them as early as day 6 (Fig.2D). Notably, some elncRNAs affected resistant cell survival only during drug exposure (Fig.2D).

The distribution of β scores in parental and resistant cells exposed to ibrutinib showed concordance among many transcripts modulating the drug response, especially among lincRNAs (Fig. 2E, left). However, some elncRNAs peculiarly affected ibrutinib efficacy in resistant or parental cells (Fig. 2E, right). Noteworthy, stable silencing of AC011294.3 (ADARreg) and RP11-96H19.1 (referred to lncPCED1b by LNCipedia, or SLC38A4-AS1 by GENCODE V49 due to its proximity to these genes (Fig.S2A)) was sufficient to revert BCR dependency in resistant cells (Fig. 2E, right).

### ADARreg and lncPCED1b affect the sensitivity to ibrutinib

To validate the CRISPRi screen results, we transduced VL51-Ibru cells with constructs expressing mCherry and pgRNAs targeting the ADARreg or lncPCED1b TSS (Fig.S2A, B) and GFP-tagged dCas9ZIM3. As a negative control, we used pgRNAs targeting the TSS of KLK12, a gene not expressed in this model. After two weeks, cells expressing both fluorescent markers were sorted and analyzed. Knockdown of ADARreg or lncPCED1b increased sensitivity to BTK inhibition in ibrutinib-resistant cells (Fig. S2C). In contrast, parental cells with lncPCED1b knockdown did not show this phenotype (Fig. S2C). Notably, in ibrutinib-resistant cells, the phenotype remained stable over time and was maintained in monoclonal lines generated weeks after selection of the polyclonal population (Fig. 2F). These findings highlight the essential role of ADARreg and lncPCED1b in sustaining BCR blockade in ibrutinib-nonresponsive cells and underscore the value of these stable models for dissecting their mechanisms of action.

### ADARreg modulates BTK inhibition by regulating ADAR2 activity

To generalize the biological relevance of ADARreg beyond the specific *in vitro* model we used for discovery, we examined ADARreg expression in B-cell lymphoma clinical datasets in which we could quantify the expression of ADARreg, and we found correlation with poorer overall survival in patients with mantle cell lymphoma (MCL) and with germinal center-like diffuse large B cell lymphoma (GCB_DLBCL) (Fig. 3A).

**Figure 3.**
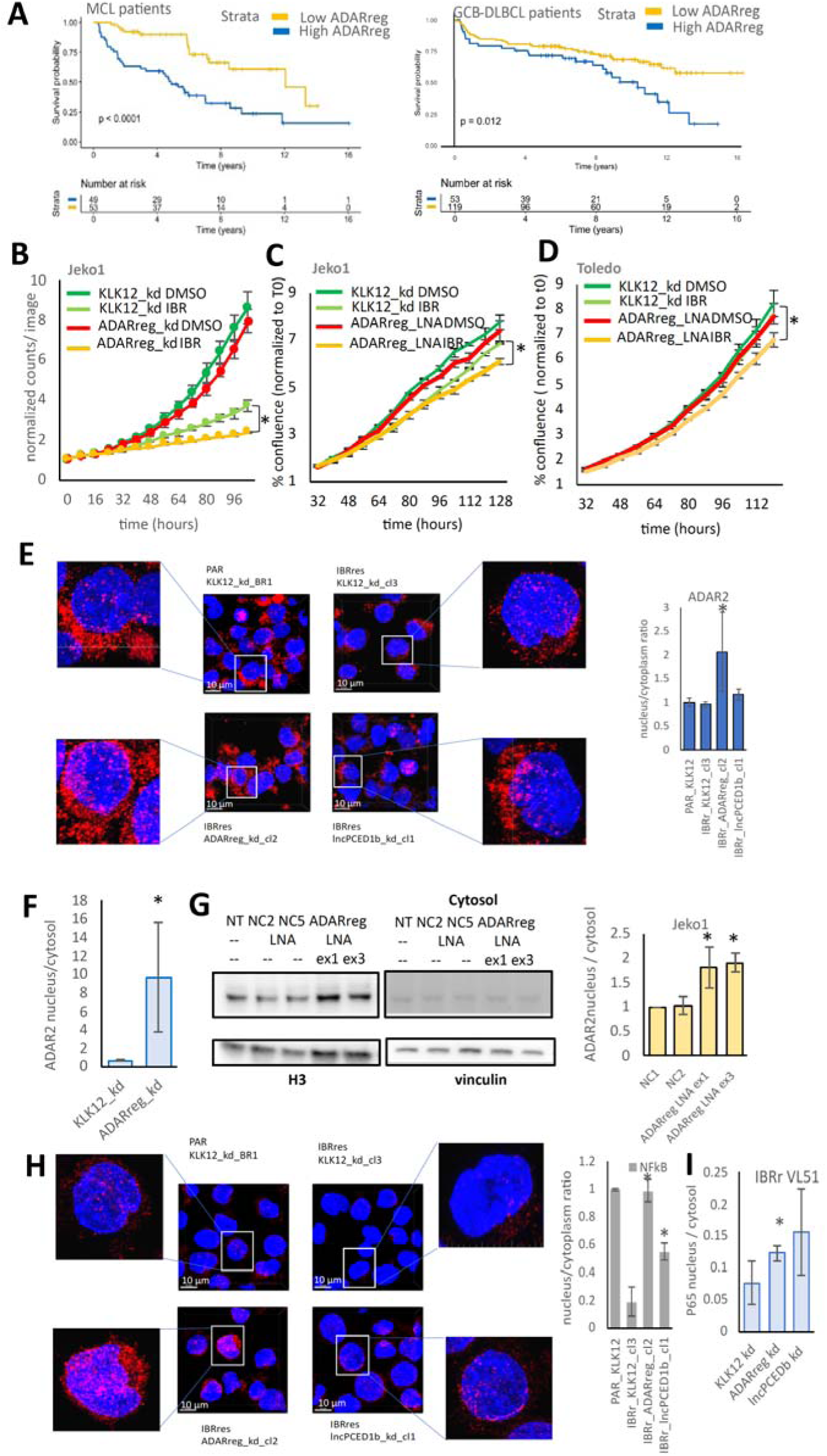
ADARreg modulates BTK inhibition response by regulating ADAR2 activity. (A) Kaplan–Meier survival of MCL (n= 102) or GCB-DLBCL (n= 172) patients stratified by high or low ADARreg expression. (B) Growth curves of Jeko1 KLK12 or ADARreg-kd cells under vehicle or ibrutinib. (C and D) Growth curve of Jeko1 (C) or Toledo (D) exposed to ibrutinib or vehicle, 4 days after ADARreg LNA or control electroporation. (E) ADAR2 immunofluorescence in parental VL51, ibrutinib-resistant KLK12-kd, ADARreg-kd, or lncPCED1b-kd cells. Representative images (left) and nuclear/cytoplasmic quantification of ADAR2 integrated density (right). DAPI-stained nuclei, blue, Alexa647-conjugated antibody-stained ADAR2, red. Close-up from each picture shows a single-cell level magnification. (F) Nuclear/cytosolic ADAR2 protein levels in ibrutinib-resistant KLK12-kd and ADARreg-kd VL51 clones. Average of three independent monoclonal cell lines per group. (G) Nuclear/cytosolic ADAR2 protein levels in Jeko1 7 days after ADARreg-LNA or controls, left, and relative quantification, right. H3 was used to normalize nuclear lysate and vinculin, the cytosolic. (H) NF-κB immunofluorescence in parental VL51, ibrutinib-resistant KLK12-kd, ADARreg-kd, or lncPCED1b-kd cells. Representative images (left) and nuclear/cytoplasmic quantification of NF-κB integrated density (right). DAPI-stained nuclei, blue, Alexa647-conjugated antibody-stained ADAR2, red. Close-up from each picture shows a single-cell level magnification. (I) NF-κB p65 nuclear/cytosolic protein levels across three monoclonal lines per group. (n=3, P.value<0.05)

Therefore, we tested the ADARreg function in Jeko1, a MCL line that expresses ADARreg and shows low sensitivity to ibrutinib. Stable ADARreg silencing significantly increased their sensitivity to ibrutinib (Fig.3B).

To confirm that ADARreg *per se* contributed to drug resistance, we used antisense oligonucleotides to degrade ADARreg in Jeko1 cells. Electroporation with Locked Nucleic Acids (LNAs) targeting ADARreg reduced ADARreg expression to a level comparable to the stable ADARreg-kd polyclonal population (Fig. S3A), and also the sensitivity to ibrutinib increased (Fig.3C). We reproduced the same phenotype in the Toledo cell line, which is totally resistant to ibrutinib in agreement with its GCB-DLBCL origin (Fig.3D). As further control of ADARreg-dependency of BTK inhibition resistance, we nucleofected ADARreg LNAs in HAIRM, an ibrutinib-sensitive cell line derived from MZL or hairy cell leukemia variant. HAIRM did not express ADARreg and did not exhibit changes in proliferation or drug sensitivity after LNA exposure (Fig.S3B).

On the contrary, we attempted to extend the role of lncPCED1b to other lymphoma subtypes; however, we did not observe any significant variation in BCR blockade sensitivity in either Jeko1 or Toledo cells (Fig. S3C, D).

Given ADARreg’s potential role in regulating A-to-I RNA editing ^20^, we studied ADARreg correlation to ADAR enzymes. Its expression was negatively correlated with ADAR1 (Fig. S3E) and positively correlated with ADAR2 (encoded by ADARB1) in the clinical specimens (Fig. S3F). However, ADARreg knockdown did not significantly alter ADAR1 or ADAR2 global protein expression, neither in VL51-Ibru nor in Jeko1 (Fig.S3G).

According to ADARreg predominant nuclear localization (Fig.S3H), we hypothesized its presence could modify ADAR enzymes’ subcellular localization instead of their expression. Indeed, ADARreg stable knockdown in VL51-Ibru showed increased ADAR2 nuclear translocation, by immunofluorescence (Fig.3E) and by immunoblotting after nuclear/cytoplasm fractionation (Fig. 3F, S3I). ADARreg knockdown did not affect ADAR1 subcellular distribution (Fig.S3J). We confirmed the increased ADAR2 nuclear localization also in Jeko1 after 7 days of ADARreg LNA exposure (Fig. 3G).

As confirmation of the recovered dependency on the BCR pathway of lymphoma cells after ADARreg silencing, we measured increased nuclear translocation of NF-κB by immunofluorescence (Fig. 3H) and by immunoblotting (Fig. 3I, S3I). Consistently, the phosphorylation of full-length p65 increased in resistant cells after BCR stimulation following ADARreg, but not after lncPCED1b knockdown (Fig.S3K).

### ADARreg silencing affects A-to-I editing in intronic regions

To assess changes in ADAR processivity, we compared transcript variants between parental or ibrutinib-resistant cells expressing KLK12 pgRNAs, and Ibrutinib-resistant cells with stable knockdowns of ADARreg or lncPCED1b versus control. Using total RNA-Seq from polyclonal cell lines and the RED-ML algorithm ^28^, we identified approximately 24,000 editing sites within and 13,000 outside Alu sequences across all conditions (Table S4).

Focusing on sequence-specific editing sites detected in at least three samples, 66% (n=1,857) mapped outside Alu sequences and 33% (n=952) within (Fig.4A, left). Differential editing analysis identified 195 non-Alu and 86 Alu-specific editing sites altered in the resistant cells. Comparable analyses of resistant cells with ADARreg knockdown (85 non-Alu; 14 Alu) or lncPCED1b knockdown (69 non-Alu; 18 Alu) showed that elncRNA silencing predominantly affected editing at non-Alu sites (Fig.4A, right).

**Figure 4.**
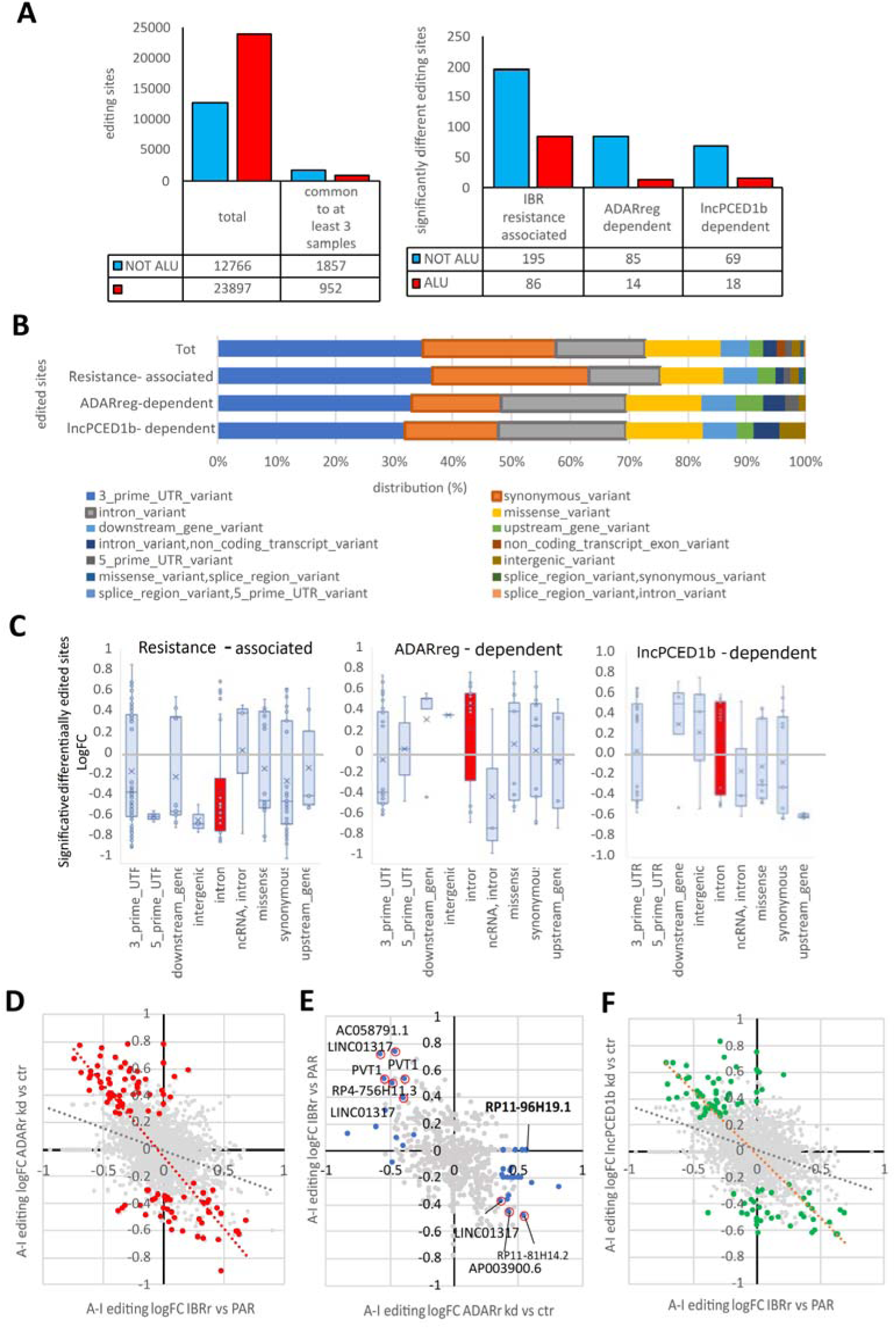
ADARreg silencing alters A-to-I editing in introns. (A) Editing sites inside Alu (red) or non-Alu (blue) regions (left), and significant differentially edited sites in ibrutinib-resistant versus parental, or ibrutinib-resistant ADARreg-kd or lncPCED1b-kd versus controls (right)(p-value < 0.05, Limma test). (B) Genomic distribution of total and differentially edited sites in ibrutinib-resistant vs parental, or ibrutinib-resistant ADARreg-kd or lncPCED1b-kd vs controls (p-value < 0.05, Limma test). (C) LogFC of non-Alu editing sites across genomic categories in ibrutinib-resistant versus parental (left) and in ADARreg-kd (middle) or lncPCED1b-kd (right) versus control (pvalue< 0.05, Limma test, mean and outlier values are shown). (D) Correlation of editing site logFC in ibrutinib-resistant vs parental and ADARreg-kd vs control; significant sites are highlighted. The regression line represents Pearson correlation (p-value < 0.05, Limma test). (E) Correlation of ncRNA editing site logFC in ADARreg-kd vs control and ibrutinib-resistant vs parental; significantly affected sites (blue) or those shared with lncPCED1b-kd (red circled) are marked (p-value < 0.05, Limma test). (F) Correlation of editing site logFC in ibrutinib-resistant versus parental and lncPCED1b-kd versus control; significant sites are highlighted. The regression line represents Pearson correlation (p-value < 0.05, Limma test).

We next examined the genomic distribution of editing sites. Approximately 30% localized to 3’UTR, 20% to exons (mostly synonymous variants), 15% to exonic missense positions, and 17% to introns (Fig.4B). ADARreg and lncPCED1b knockdown selectively altered intronic editing (Fig.4B).

In Ibrutinib-resistant cells, intronic editing was largely lost compared to parental cells (Fig.4C, left). ADARreg knockdown reversed this trend, increasing intronic editing in protein-coding genes while decreasing it in noncoding genes (Fig.4C, middle). lncPCED1b knockdown produced a similar increase in intronic editing (Fig.4C, right). Many of the editing sites restored by ADARreg knockdown corresponded to editing events originally present in parental cells but lost in resistant cells (Fig.4D). Transcripts containing differentially edited sites were enriched for immune- and cancer-related pathways (Fig.S4A).

Altogether, these findings support a role for ADARreg, and partially lncPCED1b, in redirecting not-Alu RNA editing towards introns of specific transcripts in BCR-independent lymphoma cells.

### ADARreg affects the editing of ncRNAs involved in ibrutinib sensitivity

We wondered if the observed RNA editing changes were specifically associated with ADARreg and if the activity was mediated by other lncRNAs, like lncPCED1b. Several elncRNAs and lincRNAs were differentially edited in resistant versus parental cells (Fig.4E). Notably, 37 sites showed significant differential editing upon ADARreg loss, most of which were more edited than the controls (Fig.4E). lncPCED1b (RP11-96H19.1) was the third most edited ncRNA in ADARreg knockdown cells (Fig.4E), and its knockdown partially restored the parental editome in resistant cells (Fig.4F). To further validate this, we analyzed parental cells with stable knockdowns of lncPCED1b or LINC00511, a known oncogene not associated with RNA editing. Editing sites modified upon LINC00511 knockdown showed a weaker correlation with resistance-associated sites than those affected by ADARreg knockdown (Fig. S4B–D). In contrast, lncPCED1b knockdown did not comparably reshape the editome of VL51 cells, in which ADARreg did not affect ibrutinib resistance (Fig.S4E). Furthermore, intronic editing events after lncPCED1b or LINC00511 knockdown were less pronounced in parental cells than in resistant ones (Fig.S4F-G).

### ADARreg and lncPCED1b maintain the BCR inactivation by altering RNA homeostasis

Transcriptomic analysis of cells after ADARreg or lncPCED1b knockdown revealed significant enrichment of genesets related to the proteasome, spliceosome, and RNA degradation among the transcripts more expressed in resistant compared to parental cells (Table S5, Fig.5A). Inhibiting these elncRNAs, we also reversed genesets normally suppressed by BTK inhibition, in resistant cells, confirming the reactivation of BCR dependency (Fig.5A). ADARreg and lncPCED1b inhibition reverted the SMZL signature associated with ibrutinib-resistance ^21^, confirming their role in maintaining the resistant phenotype (Fig.5A). Furthermore, resistant cells lost the differentiation features of MZ B cells (downregulation of genes upregulated in nodal MZL vs lymph node), typically driven by BCR/NF-kB signaling pathway, adhesion and interaction genes, like integrins and CD82, and cytokine receptors or antigen-presenting molecules, like CD74. ADARreg and lncPCED1b knockdown markedly reconstituted their dependence on microenvironment-driven signaling (Fig.5A). For instance, resistant cells had lower expression of genes involved in interferon response compared to parental cells. Still, ADARreg and lncPCED1b silencing reactivated immune responses through PRR signaling (Fig.5A, Table S6). Specifically, ADARreg knockdown restored the expression of interferon-stimulated genes (ISGs) such as IFIH1 and DHX58, critical in the RIG-I-like receptor (RLR) pathway that induces interferon production after detecting unedited dsRNA, normally suppressed by ADAR editing ^29^ (Fig.5B).

**Figure 5.**
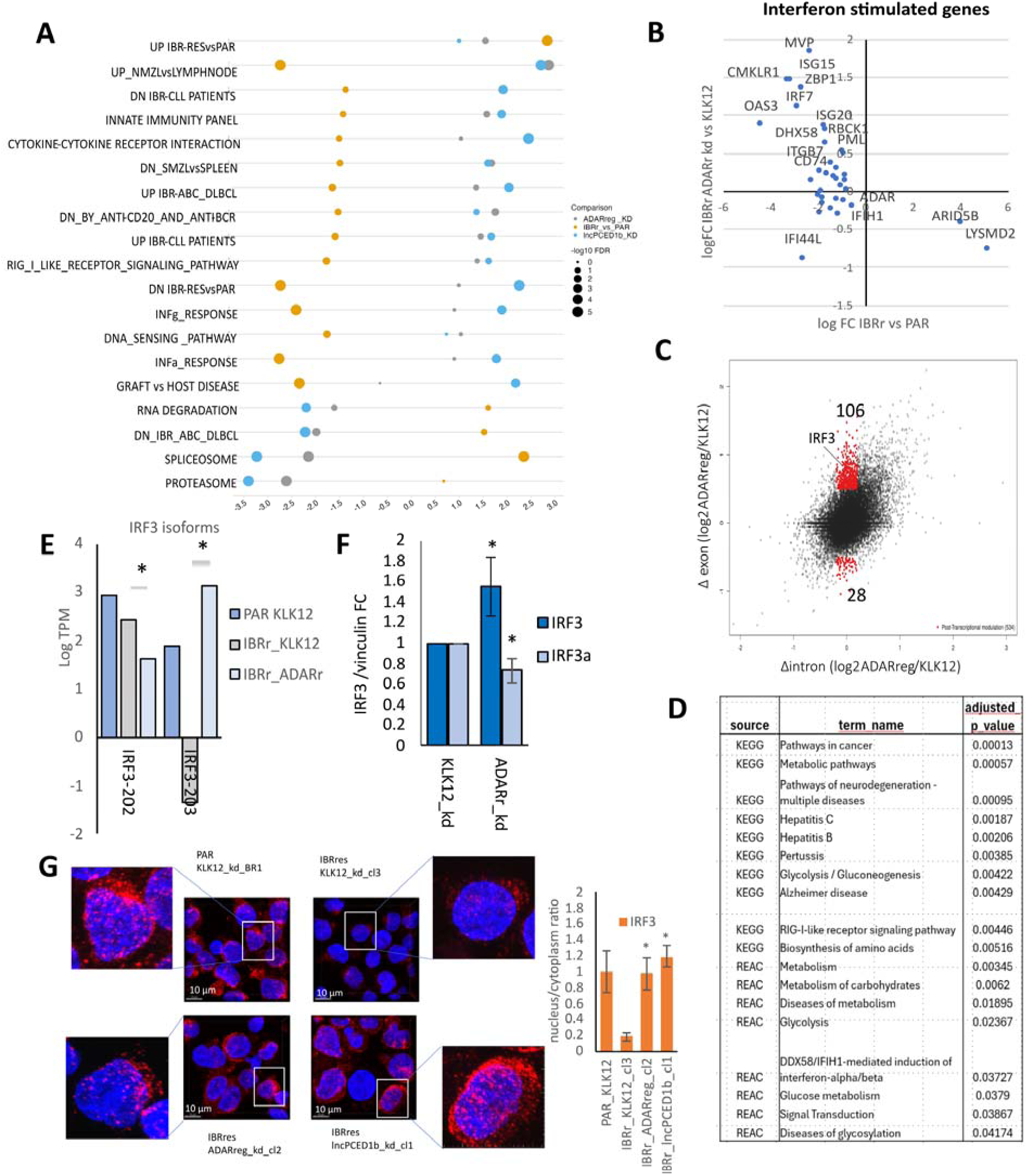
ADARreg modulates IRF3 activity. (A) GSEA of ibrutinib-resistant versus parental, ibrutinib-resistant ADARreg-kd or lncPCED1b-kd versus control. Normalized enrichment score and significance are represented. (B) Scatter plot of interferon-stimulated gene logFC in ibrutinib-resistant vs parental and ibrutinib-resistant ADARreg-kd vs control. (C) Changes in exonic (Δexon) versus intronic (Δintron) reads per transcript in ibrutinib-resistant ADARreg-kd vs control; post-transcriptionally modulated transcripts (Δexon/Δintron≠1) are highlighted (P-value<0.05). (D) gProfiler gene ontology analysis of post-transcriptionally modulated transcripts. (E) IRF3 isoform abundance in parental, ibrutinib-resistant control, and ibrutinib-resistant ADARreg-kd cells (*FDR < 0.05). (F) IRF3/IRF3a protein quantification in ibrutinib-resistant control and ADARreg-kd cells (n=3, * P <0.05). (G) Left, IRF3 immunofluorescence in parental, ibrutinib-resistant KLK12-kd, ADARreg-kd, or lncPCED1b-kd cells. DAPI-stained nuclei, blue; Alexa647-conjugated antibody-stained IRF3, red. Close-up from each picture shows a single-cell-level magnification. Right, nuclei/cytoplasm ratio of IRF3 integrated density (n=3, * P <0.05).

### ADARreg modulates IRF3 activity

Since ADARreg and lncPCED1b knockdown repressed many enzymes involved in RNA degradation (Fig. 5A), we assessed post-transcriptional modulation of transcript abundance using exon-intron split analysis (EISA). Cells with ADARreg or lncPCED1b knockdown exhibited increased transcript stability, particularly ADARreg-kd cells (106 stabilized transcripts compared to 64 for lncPCED1b) (Fig.5C, Fig.S5B, Table S7). The stabilized transcripts were involved in metabolic pathways and response to viral infection (Fig. 5D), while lncPCED1b knockdown primarily stabilized transcripts encoding proteasome components (Fig.S5B). IRF3, a key transcription factor in immune response, was post-transcriptionally regulated by ADARreg (Fig.5C). ADARreg-kd stabilized the active IRF3 isoform (IRF3-203), while destabilizing the dominant-negative IRF3a (IRF3-202) (Fig.5E, S5C) ^30^. This shift in isoform balance was confirmed at the protein level (Fig.5F, S5D), and immunofluorescence demonstrated increased nuclear localization of IRF3 after ADARreg knockdown (Fig.5G). In resistant cells, ADARreg expression reduced all IRF3 transcripts, with the active IRF3-203 isoform completely depleted (Fig.5E, S5C). After ADARreg knockdown, IRF3 target genes, typically downregulated in resistant cells, were partially reactivated (Fig.S5E).

### ADARreg alters transcript usage

Given the massive alteration of the splicing machinery after elncRNA knockdown (Fig.5A), we next investigated whether ADARreg depletion broadly affected transcript usage. Transcriptome analysis showed that ADARreg knockdown induced widespread alternative transcript usage, affecting 1,536 genes, while lncPCED1b knockdown only 469 genes (Fig.S6A, Table S8). Many of these isoform changes occurred in genes with intronic editing, with ADARreg affecting 160 and lncPCED1b 63 genes (Fig.S6A). Variations in 3’UTR editing were rarely associated with isoform changes (Fig.S6A). Some genes with sequence-specific editing, ENO1, RPS3, CLEC2D, MFN1, and AUTS2, displayed isoform switching after ADARreg knockdown, with RPS3 similarly affected by lncPCED1b knockdown (Fig.S6A).

To elucidate how ADARreg regulates isoform abundance and how this is linked to intronic editing, we performed direct RNA sequencing (DRS). This long-read sequencing approach can capture native, full-length RNA molecules, including their modifications ^31^. We precisely quantified isoform switching in ADARreg-kd versus parental cells. Nuclear and cytosolic fractions were analyzed separately to distinguish ADARreg effects on nuclear regulatory RNAs (such as elncRNAs) from those on mature cytosolic transcripts. Only full-length reads were considered for downstream analyses.

In both nuclear and cytosolic compartments, a large fraction of genes produced multiple isoforms, 30.4% in the nucleus and 40.8% in the cytosol (Fig.6A). In the nucleus, the highest fraction of genes (41.3%) transcribed noncoding isoforms, likely with regulatory function, while in the cytosol, almost the 30% produced multiple protein isoforms, indicating broad functional diversity (Fig.6A). Among nuclear noncoding transcripts, retained intron isoforms were the most abundant, followed by processed pseudogenes and lincRNAs (Fig.6B, top). Retained intron transcripts originated from both coding and noncoding genes, with a similar frequency, but the latter produced generally longer transcripts (Fig.S6B). A minor fraction of the noncoding transcripts was exported to the cytosol (Fig.6B, bottom).

**Figure 6.**
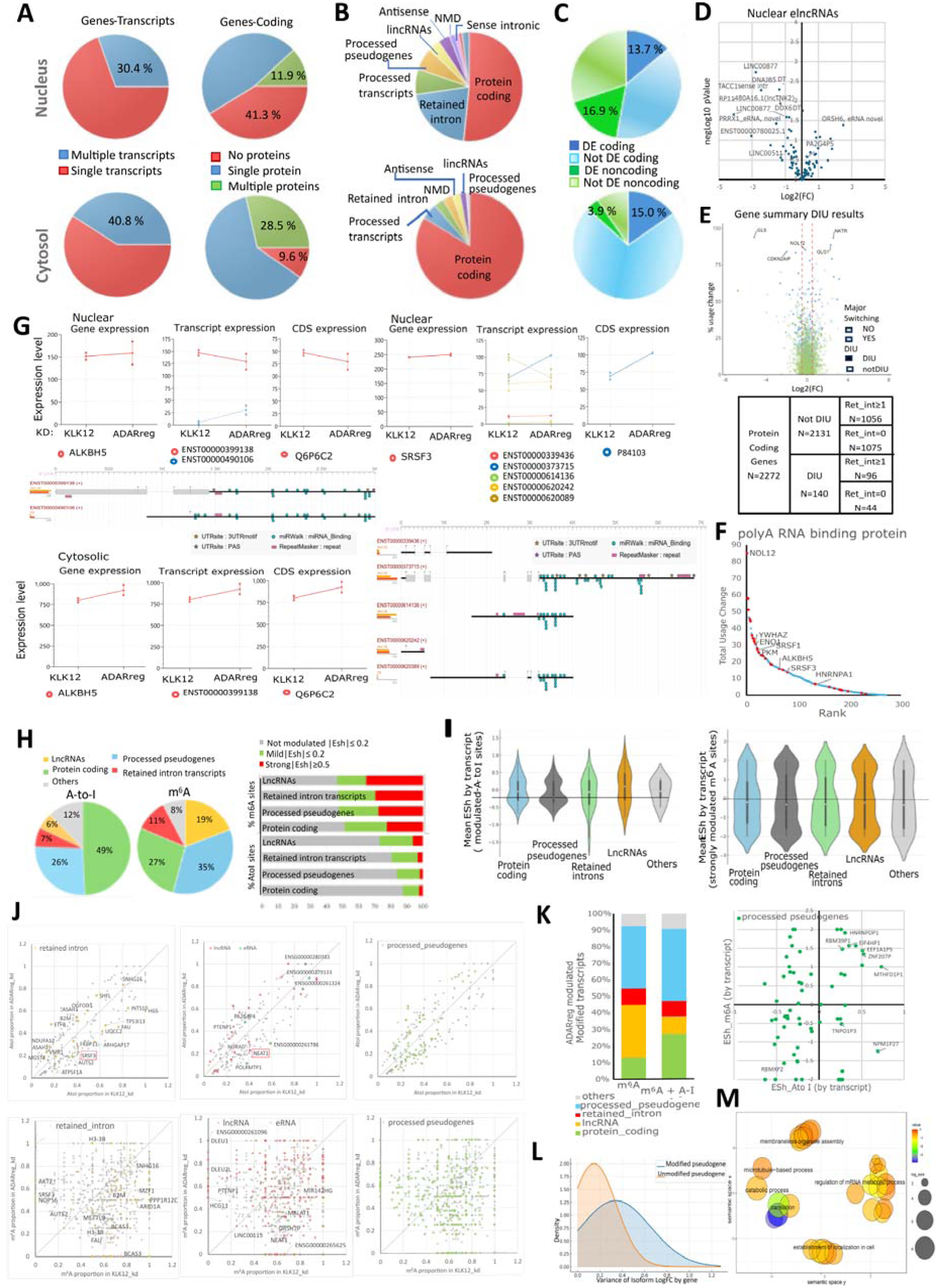
ADARreg regulates RNA homeostasis at multiple levels. DRSeq full-length RNA profiling of nuclear and cytosolic transcripts in VL51-Ibru ADARreg-kd and controls. (A) Proportion of genes producing single/multiple transcripts and coding for zero/one/multiple isoforms in the nucleus (top) or in the cytosol (bottom). (B) Nuclear (top) or cytosolic (bottom) transcript biotype distribution. (C) Nuclear (top) or cytosolic (bottom) differentially expressed coding and noncoding transcripts (FDR<0.05). (D) Volcano plot of nuclear elncRNAs. Statistically significant hits (FDR < 0.05) are labeled. (E) Top, isoform usage change versus gene expression logFC between ADARreg-kd and control in the nucleus. Top genes with major isoform switches (blue) are labeled. Bottom, numbers of protein-coding genes subject to DIU, involving a retained intron isoform, in the nucleus. (F) Ranking of isoform usage changes in polyA-RNA-binding proteins, highlighting m^6^A RNA methylation regulators. (G) Top, gene, transcript, and protein-level expression of ALKBH5 and SRSF3 in ADARreg-kd and control in the nucleus (squares, replicate expression; points, mean replicate expression). Middle, tappAS graphical representation of transcript-level annotations. (Grey boxes, exons; dotted lines, spliced introns; black line noncoding retained sequences). Bottom, ALKBH5, gene, transcript, and protein-level expression in the cytosol. (H) A-to-I and m^6^A site distribution by biotype of nuclear transcripts detected by DRS full-length-read, left, and by effect size, in the comparison ADARreg-kd versus control (not modulated (|ESh|<0.2), mild (0.2<|ESh|<0.5), strong (|ESh|>0.5)) (I) Aggregated effect sizes (ESh mean of modulated sites by transcript) for A-to-I, left, and m^6^A, right, across transcript biotypes. (J) Proportion of A-to-I (top) and m^6^A (bottom) significantly modulated sites, aggregated by transcript, in control and ADARreg-kd samples. Retained intron transcripts, lncRNAs, and pseudogenes are highlighted. (K) Top, distribution of nuclear transcripts carrying ADARreg-dependent m^6^A ± A-to-I changes by transcript biotype; bottom, scatter plot of A-to-I and m^6^A ESh values of nuclear processed pseudogenes. (L) Variance distribution of the cytosolic isoform logFC by parent gene, paired with differentially edited/methylated nuclear pseudogenes after ADARreg-kd (blue), or paired with only differentially methylated nuclear pseudogenes (orange). Mean, µ, and standard deviation, W, of the variance are indicated. (M) Semantic clustering of enriched gene ontology terms in parent genes of m^6^A/A-to-I-modulated pseudogenes

ADARreg knockdown markedly altered the abundance of nuclear transcripts (Fig.S6C-D), primarily of ncRNA (Fig.6C, top), and affected approximately 15% of protein-coding transcripts, consistently in both the nucleus and the cytosol (Fig.6C, bottom). Among 100 full-length elncRNAs detected in the nucleus, several were downregulated by ADARreg knockdown (Fig.6D). In the cytosol, 161 elncRNAs, were identified, equally up- and downregulated, suggesting indirect effects (Fig.S6E).

Differential isoform usage (DIU) was quantified by TappAS algorithm ^32^, comparing global expression fold changes with isoform redistribution. Globally, we noticed stronger DIU in nuclear than in cytosolic transcripts (Fig.6E, S6F), although some genes, such as BAG1, displayed clear isoform switches at the mature mRNA level (Fig.S6G). Nuclear-cytosolic comparisons suggested that ADARreg also regulated mRNA export. For example, CDKN2AIP expression decreased globally after ADARreg knockdown, but one truncated isoform (J3KNE1) was retained in the nucleus, while the main isoform (Q9NXV6), although downregulated, remained predominant in the cytoplasm (Fig.S6H).

Among 2,272 genes, corresponding to 5,293 protein-coding isoforms detected in the nucleus, 140 were differentially expressed. Seventy-five % of them (n=96) transcribed at least one retained intron transcript (Fig.6E, Table S9). They were enriched for genes coding for polyA RNA binding proteins (n=25) (Fig.6F, S6I), notably ALKBH5 and SRSF3 (Fig.6G). ALKBH5 encodes a m^6^A eraser involved in BCR signaling^33^ and in the tumor microenvironment ^34^, whereas SRSF3 is a splicing factor, interactor of m^6^A reader YTHD1 ^35^ and already reported to regulate lncRNA splicing ^36^. Both genes exhibited nuclear retained-intron isoforms (Fig.6G). The nuclear accumulation of the ALKBH5 retained intron isoform coincided with augmented cytosolic export of the protein-coding form (Fig.6G). These data suggest that ADARreg modulates the nuclear abundance of retained intron transcripts, thereby post-transcriptionally regulating genes involved in RNA homeostasis.

### ADARreg regulates the editing of nuclear intron-retaining transcripts

To uncover how ADARreg contributes to RNA homeostasis, we analyzed the RNA modification events at the isoform level. We computed the A-to-I editing variation in the samples, linking the inosine modification to a specific RNA isoform only if supported by full-length reads. 49% of the nuclear editing events occurred in protein-coding isoforms, 26% in processed pseudogenes, 7% in retained introns and 6% in lncRNAs (Fig.6H, left). These percentages partially reflected the abundance of each biotype. However, using Cohen’s index (ESh) to quantify effect size, ADARreg knockdown showed mild (0.2<|ESh|<0.5) or even strong (|ESh|>0.5) impact on A-to-I editing, particularly in lncRNAs (e.g. NEAT1) and retained intron transcripts (e.g. SRSF3) (Fig.6H, right, Fig.6I-J). Thus, editing changes within introns, previously described in polyclonal elncRNA-silenced cells (Fig.4C), primarily reflected altered nuclear editing of retained-intron transcripts (Fig.6I-J, top).

In contrast, cytoplasmic editing events were almost exclusively in protein-coding isoforms, though the few lncRNA-associated sites (0.6%) were strongly ADARreg-dependent (Fig.S6J).

### ADARreg regulates RNA homeostasis also by means of m⁶A RNA methylation

The modulation of nuclear abundance of retained intron transcripts and mRNAs encoded by ALKBH5 and SRSF3, or ENO1 and PKM, well-recognized m⁶A targets^37,38^, suggested that ADARreg might also influence RNA methylation. Applying isoform-resolved m⁶A quantification, we found that 35% of nuclear m⁶A peaks occurred in processed pseudogenes (Fig.6H), consistent with the well-documented role for m⁶A-coupled decay of those transcripts ^39^. ADARreg knockdown significantly altered m⁶A abundance in 30-40% of sites across major transcript biotypes, including protein-coding, pseudogene, retained-intron, and lncRNA isoforms (Fig.6H). Unlike A-to-I editing, strongly m⁶A-regulated sites were similarly distributed across transcript classes (Fig.6I, right). Consistent with increased cytosolic ALKBH5 mRNA, overall nuclear m⁶A levels tended to decrease upon ADARreg knockdown (Fig.6I–J).

We investigated whether variations in A-to-I editing could affect the m⁶A abundance at specific targets. Comparing nuclear transcripts with exclusively m⁶A modulation and transcripts where both m⁶A and A-to-I significantly changed after ADARreg knockdown revealed that both marks were significantly different in protein-coding genes and pseudogenes (Fig.S6K, 6K). The increased A-to-I events in pseudogenes were associated with an increase in m⁶A (Fig.6K, right). To investigate the role of RNA editing in the context of nuclear pseudogenes, we compared the differential expression of cytosolic isoforms of paired parent genes, specifically when the pseudogene was modulated in terms of m⁶A and A-to-I editing, or m⁶A only. Parent genes of edited pseudogenes displayed greater isoform variability (logFC variance mean, µ = 0.34) than those paired with only methylated pseudogenes (logFC variance mean, µ = 0.146) (Fig.6L). Parent genes paired with edited pseudogenes were primarily involved in RNA homeostasis, controlling mRNA metabolic processes and localization, specifically NPM1 and HNRNPD (Fig. 6M).

For nuclear protein-coding transcripts bearing both modifications, increased editing was mainly associated with reduced m⁶A. These transcripts were predominantly mono-exonic (e.g., histone genes), typically more methylated than spliced RNAs ^40,41^. The A-to-I editing in these transcripts could induce their demethylation and influence stability (Fig. S6K).

### ADARreg affects protein-coding isoforms involved in immune regulation

To investigate the effects of the observed transcriptional and post-transcriptional changes, we performed a functional enrichment analysis of the protein-coding isoforms expressed in the cytosol, using the TappAS algorithm ^32^. We identified four clusters of enriched gene ontology terms: cluster 1, related to chemokine-mediated signaling pathway (e.g., CCR5 chemokine receptor binding); cluster 2 and 3, related to the negative regulation of immune effector process and of interferon-gamma production; cluster 4, related to the positive regulation of calcium ion transport (Fig.7A). Key upregulated transcripts included CD38, a marker of BCR activation ^42^, and XBP1, which regulates plasma cell differentiation ^43^. Conversely, TNFSF4, CCL3, CCL4, CCL5, XCL1, and IL23A were downregulated, indicating a potentially impaired recruitment of a protective immune niche ^44^. Reduced CEACAM1, which modulates BCR signaling ^45^ and NK cell activity ^46^, further supports immune suppression by ADARreg.

**Figure 7.**
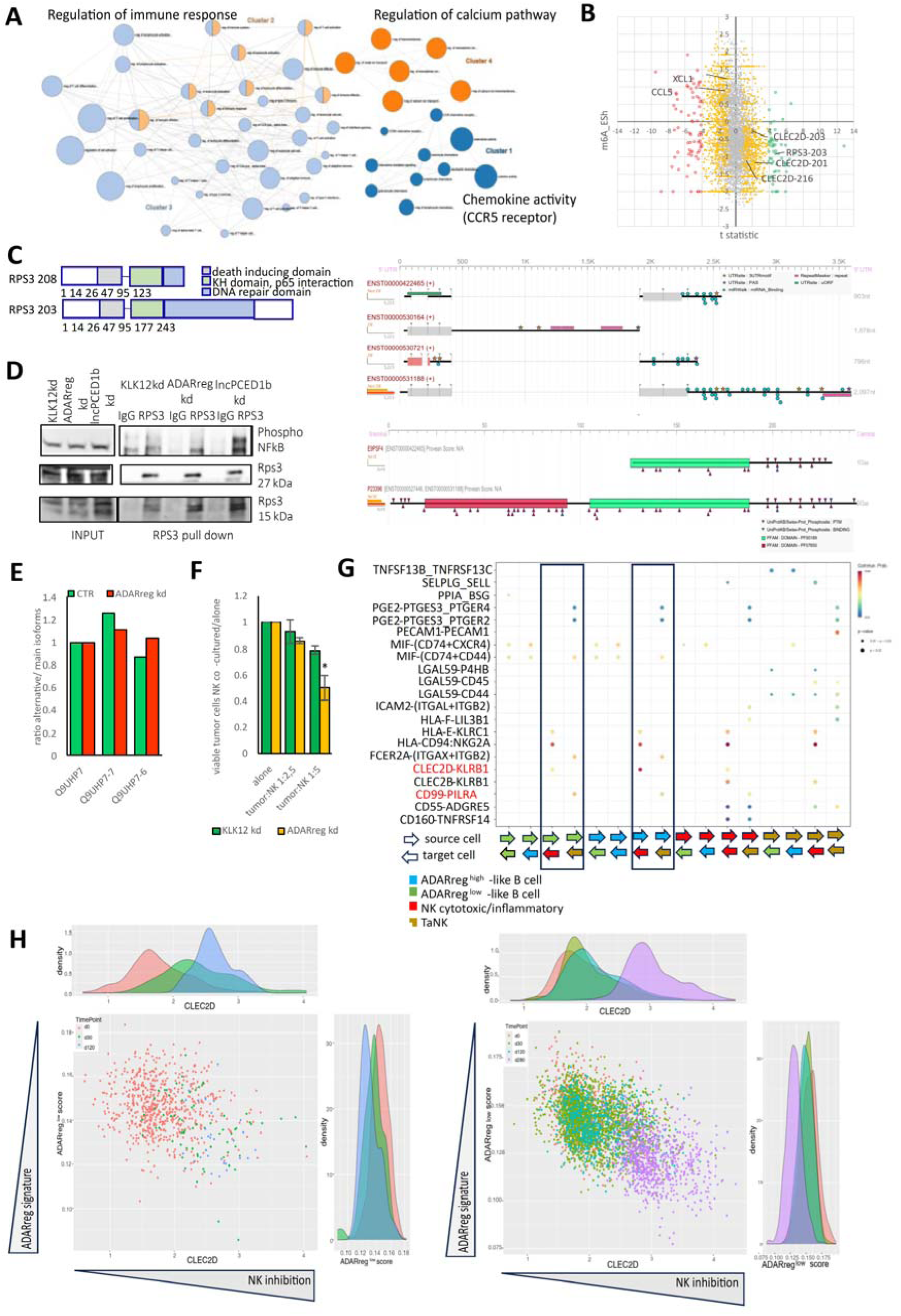
ADARreg suppresses NK-cell cytotoxicity. (A) GO cluster analysis of differentially expressed cytosolic protein-coding transcripts in ibrutinib-resistant ADARreg-kd vs control cells. Pie chart area represents DE regulation measured by relative - log10(p value) of the enrichment test, and node size represents the number of genes annotated for each GO term. (B) Correlation of t-statistic for ADARreg-kd versus control and transcript-aggregated m^6^A ESh in cytosolic protein-coding transcripts. Strongly significantly downregulated (red), upregulated (green) or mildly modulated (yellow) transcripts are highlighted. (C) Left, domain structures of rps3 isoforms encoded by RPS3-203/208. Right, TappAS graphical representation of transcript-level and protein-level annotations of RPS3. Grey boxes, exons; dotted line, spliced introns; black line, noncoding retained sequences; red and green boxes, protein domains. (D) Co-immunoprecipitation of phosphorylated p65 with long and short rps3 isoforms in ibrutinib-resistant ADARreg-kd, lncPCED1b-kd, or control cells. (E) Expression of CLEC2D protein-coding isoforms by DRS in ibrutinib-resistant ADARreg-kd versus control. (F) Viable GFP⁺-mCherry⁺ tumor cells after coculture with donor NK cells for 7 days, normalized to tumor-alone controls (n=3,* P <0.05). (G) CellChat-derived communication networks between ADARreg-high or -low B cells and NK subsets in an ibrutinib slow-responder CLL patient. (H) Single-cell expression of CLEC2D and the ADARreg signature in B cells from fast or slow ibrutinib responders at pretreatment and during therapy.

To investigate the direct transcriptional effect of ADARreg knockdown, we correlated the modulation of expression and methylation of each isoform after ADARreg silencing. Coherent with the observations in the nucleus, we found that the most significantly modulated isoforms lost m^6^A marks and were upregulated, including the isoforms of the genes coding for the ribosomal protein S3 (RPS3) and CLEC2D, the ligand for the NK/T-cell receptor CD161 ^47,48^ (Fig.7B), which had already been robustly characterized by an alternative transcriptional mechanism and intronic editing (Fig. S6A).

The most downregulated transcripts, encoding CCL5 and XCL1, were enriched for m^6^A marks after ADARreg knockdown, consistent with enhanced transcript decay (Fig.7B).

DRS also validated IRF3 isoform modulation: the longer, active isoform was stabilized after ADARreg knockdown, likely via competition with a retained-intron isoform sharing its 3W UTR miRNA sites (Fig. S7A, Fig.4C). These findings support that the effects on RNA stability due to ADARreg depletion are likely mediated by a precise deposition of m^6^A marks on selected isoforms in an ADARreg-dependent manner.

### ADARreg modulates the RPS3 and NF-κB interaction

To demonstrate the impact of the ADARreg-regulated isoform on BCR blockade in lymphoma cells, we investigate the role of *rpS3*. The latter is a multifunctional protein involved in translation, DNA repair, apoptosis, and immune regulation, by interaction with p65 to activate specific genes ^49^. RPS3 has different domains responsible for apoptosis and DNA damage (Fig.7C)^50^. ADARreg and lncPCED1b knockdown restored expression of the RPS3-208 isoform (ENST00000527273), which was low in resistant cells (Fig. S7B). This isoform lacks the DNA repair domain but retains the apoptosis-inducing and p65-interacting domains (Fig.7C, left).

Coimmunoprecipitation confirmed that ADARreg or lncPCED1b depletion increased N-terminal truncated rps3 and the interaction of rps3 with the phosphorylated form of p65 in the nucleus (Fig. 7D). DRS confirmed multiple RPS3 isoforms, including the truncated variant retaining the p65-binding domain and harboring a long 3W UTR derived from intron 4–5 retention (Fig. 7C, right). Altogether, these data suggest that RPS3 isoform regulation is a key mechanism linking ADARreg to NF-κB-driven immune reactivation.

### ADARreg inhibits NK cell-mediated cytotoxicity

Given the modulation of the interferon response through IRF3, the RPS3-driven p65 activity, and the chemokine-related pathways, we hypothesized that ADARreg silencing could enhance lymphoma immunogenicity. ADARreg knockdown reduced CEACAM1 and altered CLEC2D (LLT1) isoform usage (Fig. S7C, left). DRS detected multiple CLEC2D isoforms (Fig. S7C, right) with ADARreg knockdown favoring a shorter, less methylated isoform (Fig. 7B, E, S7D), suggesting functional modulation of NK-cell interaction.

Co-culturing ADARreg knockdown VL51 cells with NK cells from a healthy donor resulted in significantly higher NK cell-mediated killing compared to control cells (Fig. 7F, S8A). Additionally, NK cells survived longer in co-cultures with ADARreg knockdown cells (Fig. S8B).

To assess clinical relevance, we analyzed single-cell RNA-seq data from chronic lymphocytic leukemia (CLL) patients undergoing ibrutinib therapy ^51^. Using genes altered by stable ADARreg-kd as a surrogate for ADARreg level (undetectable in polyA datasets), we defined an ADARreg^low^-score to compare CLL cells from two fast and two slow responders (Fig.S8C). Transcriptomic profiles of surviving cells after treatment resembled the SMZL ADARreg^high^-VL51-Ibru model (Fig. S8C). Similar cells were also more abundant at the pretreatment time point in slow responders (Fig.S8C). We analyzed NK cells in untreated samples, annotating them as cytotoxic, inflammatory, or tumor-associated NK (TANK)-cells (Fig.S8D) ^52^, and the tumor cells, on the basis of ADARreg signature, as ADARreg^high^-like or ADARreg^low^-like B cells. Therefore, we analyzed cell-cell communication using CellChat (Fig.7G,S8E). At pretreatment, in a slow responder, ADARreg^high^-like B cells increased CLEC2D-CD161 interactions with active NKs, compared to ADARreg^low^-like B cells (Fig.7G). Conversely, ADARreg^low^-like B-cells interacted more with TANK-cells through CD99-PILRA signaling, recently associated with favorable immunotherapy outcomes^53^ (Fig.7G). No differences in cell communication between ADARreg ^high/low^-like B- and NK-cells were significant in the fast responder (Fig.S8E). Tumor cells persisting after treatment had higher CLEC2D expression and ADARreg signature in both patients (Fig.7H). However, those cells were more abundant in slow than in fast responders (Fig.7H). In summary, ADARreg modulated CLEC2D isoform balance, favoring NK inhibition in ibrutinib-resistant cells in vitro. A transcriptional program, like the one controlled by ADARreg, linked high CLEC2D expression to ibrutinib resistance in CLL patients. Finally, the tumor cells resembling the ADARreg ^high-^immune-silent VL51 interact with protumoral NK-cells through CLEC2D.

## Discussion

This study demonstrated that LOC730338, which we renamed ADARreg, plays a key role in maintaining immune suppression and refractoriness to BCR signaling regulation in lymphoma cells. We first identified its role in modulating sensitivity to the BTK inhibitor ibrutinib in an *in vitro* model of MZL with secondary BTK inhibitor resistance and subsequently extended these findings to MCL and GCB-DLBCL models, as well as CLL patients. Collectively, our results link ADARreg to the control of the inflammatory response through a complex regulatory network, based on RNA homeostasis (Fig.S9).

Although ADARreg was previously associated with RNA editing, its function remained unknown ^20^. The connection between RNA editing and immune regulation is well established: upon pathogen recognition, PRRs activate pathways such as NF-κB, MAPK, and type I interferon ^54,55^, which must be tightly controlled to prevent excessive activation^56^. ADAR1 and ADAR2 maintain immune tolerance through A-to-I RNA editing of self-dsRNA ^57^ and by modulating the transcript stability and alternative splicing of genes involved in the antiviral response ^58^. Since A-to-I editing can drastically alter RNA’s structural and coding potential ^59,60^ perturbation of this process has profound effects. Indeed, ADARreg knockdown disrupted RNA degradation and splicing machineries, highlighting its importance in maintaining RNA homeostasis.

Mechanistically, ADARreg regulates the nuclear translocation of ADAR2, a less abundant but more sequence-specific editor than ADAR1. An initial analysis of RNA editing on transcriptomic data revealed editing alteration upon ADARreg knockdown, particularly outside repetitive Alu sequences, consistent with an ADAR2-dependent mechanism ^29,61^. Notably, ibrutinib-resistant cells displayed a distinct RNA editing pattern that was reversed by ADARreg knockdown. Editing changes also affected noncoding transcripts, including elncRNAs such as lncPCED1b, suggesting a link between ADARreg and epigenetic adaptation to drug resistance. Indeed, we observed altered SEs activation and elncRNA transcription in cells with acquired refractoriness to BCR activation, implicating ADARreg and lncPCED1b in the BCR pathway regulation.

Our findings suggest that ADARreg operates within a broader network of ncRNAs that are modulated by ADARreg itself and are subject to RNA modifications. Some ncRNAs, such as lncPCED1b, appear to mediate ADARreg’s transcriptomic effects. We propose that ADARreg, controlling ADAR2 localization and elncRNA expression, regulates the stability and localization of protumoral transcriptional isoforms. Consistent with this, ADAR2 has previously been reported as a potential tumor suppressor in glioma ^62^ , and alternative splicing frequently promotes pro-tumorigenic isoforms ^60,63^.

Although globally informative and practical, RNA editing estimation on total RNA-seq cannot fully resolve how A-to-I editing affects isoform-specific splicing, as multiple transcripts may overlap the same editing sites. Recent advances in chemistry and analytical tools have made DRS a powerful technology to study isoform switching and isoform-resolved RNA modifications in subcellular compartments ^31,64,65^. Using this strategy and leveraging our adapted *in silico* pipeline to enhance the single-molecule resolution of the method, we found that ADARreg regulates the A-to-I editing of nuclear-retained intron transcripts and influences the alternative expression of genes involved in RNA homeostasis. These include multiple RNA-binding proteins that regulate RNA trafficking, maturation, and post-transcriptional modifications, like the m^6^A eraser ALKBH5 and the SRSF3, interactor of the m^6^A reader YTHDC1 ^66^. Accordingly, ADARreg knockdown reduced nuclear m⁶A levels, consistent with the downregulation of ALKBH5. Interestingly, inosine modifications appeared to protect specific pseudogenes from RNA demethylation, enhancing their nuclear retention or stability. The modulation of pseudogenes, which sequester miRNAs and other binding factors in the cytosol, exposes parent genes to altered post-transcriptional regulation of their isoforms. One key target of this mechanism, NPM1, encodes nucleophosmin 1, a nucleolar protein that forms the NPM–ALK fusion in anaplastic large cell lymphoma and is deregulated in up to 30% of acute myeloid leukemias through exon 12 mutations that cause its aberrant cytoplasmic mislocalization ^67^. Given its nucleolar localization and shuttling activity, NPM1 may also influence ADAR2 trafficking; however, a direct interaction remains unproven. Notably, NPM1 also acts as a chaperone for NF-κB ^68^ and promotes immune evasion in several cancers ^69^.

The modulation of RNA nuclear retention is also linked to ADARreg through its impact on NEAT1, a nuclear lncRNA with scaffold function in organizing paraspeckles ^70^. Those nuclear bodies are required for the storage of proteins, like transcription factors, and RNA species, like A-to-I edited mRNAs and pri-miRNAs, as a survival strategy to prepare the cell for stress ^71^. NEAT1 is also a documented target of the ALKBH5 enzyme, which stabilizes it to modulate the availability of repressors and the expression of immune-stimulating chemokines ^72^. Several observations suggest that NEAT1 and paraspeckles can influence the innate immune response, although the complete mechanism remains to be fully elucidated ^73^.

Overall, our data suggest that ADARreg influences immune signaling by modulating the BCR receptor and the downstream NF-κB signaling pathway at multiple levels. Like the reported interaction between ALKBH5 and a lncRNA (treRNA1) that modulates key transcripts, including those involved in BCR signaling in DLBCL ^33^, ADARreg appears to drive a comparable mechanism. Among its downstream targets, RPS3 was the most demethylated and upregulated cytosolic transcript following ADARreg silencing. RPS3, beyond its ribosomal functions, is involved in apoptosis, DNA repair, and transcriptional regulation ^74^. It was identified as a non-Rel subunit of NF-κB responsible for activating its immunogenic signature ^49,75^. We identified two RPS3 isoforms, with a p65-interacting domain, that enhance nuclear binding of phosphorylated NF-κB upon knockdown of ADARreg or its downstream lncRNA target, lncPCED1b. Finally, BCR and cytokine receptor signaling were reactivated, restoring sensitivity to BCR blockade.

ADARreg also contributes to immune evasion by inhibiting interferon response and favoring the expression of pro-tumoral cytokines. The active IRF3 isoform is post-transcriptionally downregulated by ADARreg, which induces the expression of a retained-intron transcript that competes with the main isoform for miRNA binding. IRF3 is a downstream target of the TLR pathway triggered by pathogen infection and is crucial for the induction of interferon-stimulated genes, as well as the subsequent innate response. Consequently, ADARreg knockdown enhanced innate immune activation and increased SMZL cell susceptibility to NK cell cytotoxicity by modulating LLT1 (CLEC2D). Notably, CLEC2D expression correlated with m⁶A-dependent isoform changes. Conversely, pro-tumoral cytokines such as CCL5 and XCL1 were destabilized due to increased m⁶A, demonstrating that ADARreg does not globally alter methylation but fine-tunes epitranscriptomic marks to restore immune competence. In SMZL patients, poorer outcomes are associated with an immunosuppressive phenotype with immune checkpoint activation ^76^. These patients often exhibit KLF2 and NOTCH2 mutations, but a clear mechanism for immune escape is missing. ADARreg may be a critical factor influencing tumor-microenvironment interactions, as sustained by the worse outcome in ADARreg^high^-GCB-DLBCL patients, who predominantly exhibit “immune-cold” microenvironment signatures ^77^. This mechanism was further validated in CLL patients, where we observed a link between the selection of tumor clones resistant to BCR inhibition and the suppression of an immunological signature modulated by ADARreg. Moreover, an ADARreg^high^-signature in hematological tumors, such as MZL, CLL, and MCL, may indicate an unfavorable prognosis for patients treated with BCR signaling inhibitors. Conversely, ADARreg represents a promising therapeutic target for antisense oligonucleotide strategies to elicit spontaneous immune responses against immune-suppressive cells.

## Limitations of the study

Although our results clearly associated ADARreg with the hijacking of a complex regulatory mechanism of RNA processing and with the subversion of the tumor-microenvironment immune crosstalk, we have not yet identified how ADARreg modulates all this in the first place. The broad molecular events conditioned by ADARreg knockdown imply that it is a highly upstream node in the network that directs the epigenetic remodeling of immune-silent lymphomas. Furthermore, additional experiments will be necessary to identify the direct RNA or protein interactors of ADARreg.

## Acknowledgments

This work was partially supported by the Swiss National Science Foundation grants CRSK-3_190808, SNSF 310030_197466, and SNSF 31003A_163232/1. NM was supported by a Ph.D. Fellowship of the NCCR RNA & Disease, a National Center of Competence in Research funded by the Swiss National Science Foundation (grant numbers 182880 and 205601). SR was funded by Science Foundation Ireland under Grant number [18/CRT/6214] and in part by the EU’s Horizon 2020 research and innovation program under the Marie Sklodowska-Curie grant H2020-MSCA-COFUND-2019-945385.

## Author contributions

L.C.: conceptualization, analysis, visualization, and manuscript preparation; F.G.: investigation and analysis; S.R.: CRISPRi library design; A.R.: investigation; F.S.: investigation; A.Z.: investigation; C.T.: investigation; S.Z.: analysis; N.M.: investigation; S.A.: investigation. A.J.A.: investigation and analysis; R.G.: investigation and resources. R.J.: conceptualization and resources; F.B.: conceptualization and resources; S.N.: conceptualization, investigation, analysis, visualization, manuscript preparation, project supervision and resources.

## Disclosure of Potential Conflicts of Interest

Luciano Cascione: institutional research funds from Orion; travel grant from HTG. Alberto J. Arribas: travel grant from Astra Zeneca, consultant for PentixaPharm. Chiara Tarantelli: travel grant from iOnctura. Francesco Bertoni: institutional research funds from ADC Therapeutics, Bayer AG, BeiGene, Floratek Pharma, Helsinn, HTG Molecular Diagnostics, Ideogen AG, Idorsia Pharmaceuticals Ltd., Immagene, ImmunoGen, Menarini Ricerche, Nordic Nanovector ASA, Oncternal Therapeutics, Spexis AG; consultancy fee from BIMINI Biotech, Floratek Pharma, Helsinn, Immagene, Menarini, Vrise Therapeutics; advisory board fees to institution from Novartis; expert statements provided to HTG Molecular Diagnostics; travel grants from Amgen, AstraZeneca, iOnctura. The other Authors have nothing to disclose.

## STAR METHODS

### Cell lines, LNA transfection, and drug treatments

All cell lines were grown in RPMI supplemented with 10% heat-inactivated fetal bovine serum (FBS). The VL51 parental and VL51-Ibru were obtained as previously described ^11,78^. HEK293T cells used for lentiviral particle production were grown in DMEM supplemented with 10% FBS. Cells were maintained at 37°C in humidified incubators under 5% CO_2_. All cell lines were tested negative for Mycoplasma contamination and were verified for identity by STR profiling. LNA ADARreg1-3 and negative controls NC1-2 were bought from IDT, Integrated DNA Technologies. Sequences are reported in Supplementary Table S10. Tumor cells (200.000 per sample) were resuspended in 20 ml of SG solution (Amaxa-Lonza) with LNA (1 nmol) and electroporated using 4D Nucleofector (Amaxa-Lonza), according to the manufacturer’s instructions and incubated for 96h, before further treatment or RNA extraction. Cells were exposed to ibrutinib (Selleckchem, Houston, TX, USA) or DMSO (Sigma) for 96h.

### IgG stimulation

VL51 parental or ibrutinib-resistant cells (5 million) were collected, washed with PBS, and exposed to 20 mg/ml of IgG (Goat Fab Anti-Human IgG, UNLB) for 20 min. Cells were washed and lysed for protein extraction.

### Cell proliferation assays

Response to drug treatments was evaluated at the endpoint by seeding 100,000 cells in two sister 24-well plates, each with serial dilutions of the drug or vehicle. The mitochondrial functionality was measured by MTT (3-(4,5-dimethylthiazol-2-yl)-2,5-diphenyl tetrazolium bromide) assay at 0 or after 96 h. Cell growth curves were measured by the Incucyte instrument. In detail, stable infected cells or transient silenced cells were seeded 5,000 cells/ well for VL51, 10,000 for Jeko1, or 20,000 for HAIRM in the presence of ibrutinib (5 mM for VL51-Ibru, KLK12kd or ADARreg kd; 40 nM for Jeko1, KLK12kd or ADARreg kd, 1 mM for LNA-transfected Jeko1, or 20 nM for LNA-transfected HAIRM and followed over time. Green fluorescence was measured for VL51, while cell-by-cell and confluence analyses were performed for Jeko1 and HAIRM cells.

### ChIP-seq

The ChIP experiment followed standard protocols with an H3K27ac (Active Motif Cat# 39133, RRID:AB_2561016) H3H4me1 (Abcam Cat# ab8895, RRID:AB_306847) and H3K4me3 (Millipore Cat# 17-614, RRID:AB_11212770) antibody or control IgG (Millipore Cat# 12-371, RRID:AB_145840) as negative control. In brief, 50 million cells were resuspended and fixed with 1% methanol-free formaldehyde. Nuclei were extracted, washed and resuspended 25 million/ml in sonication buffer (10 mM Tris pH 8.0, 100 mM NaCl, 1 mM EDTA, 0.5 mM EGTA, 0.5% N-lauroylsarcosine sodium salt and fresh 0.1% Na-deoxycholate and proteinase inhibitors). Chromatin was fragmented by sonication with Covaris to an average size of 200-500 bp. A 50 μL aliquot was saved as input, whereas 650 μL (corresponding to 16 millions of nuclei) each were incubated with 10 μg antibody, overnight at 4°C. DNA-protein complex was recovered overnight at 65°C, then proteins and RNA were digested and DNA purified by PCR purification kit (Qiagen)

Library preparation from 5 ng purified chromatin was performed using NebNext Ultra II DNA Library Prep with magnetic purification beads (E7103S, New England Biolabs) and multiplex oligonucleotides (Dual Index Primers; E7600S, New England Biolabs), followed by adaptor ligation according to the manufacturer’s protocol. Quality controls of amplified chromatin fragments were performed on Bioanalyzer 2100 (Agilent Technologies) and Qubit V4 (Thermo Fisher Scientific). Next-generation sequencing was performed on a NextSeq 2000 (Illumina) using the P2 reagent kit V3 (100 cycles; Illumina). Samples were processed starting from stranded, single-ended 120bp-long sequencing reads. Raw data in Fastq format were first preprocessed to ensure suitability for downstream analysis. Adapter sequences were trimmed from the reads using trimmomatic algorithm^79^. Quality control was performed to filter out reads with low-quality bases, short lengths, or those suspected to be sequencing artifacts. Reads were mapped to the HG38 human genome using BWA alignment. Peak calling was performed using MACS (version 2) with “broadPeak” setting to identify broad regions of H3K27me3, then annotated to the nearest genes with HOMER considering both transcription start and end sites.

Raw and processed data have the accession number (pending) and were uploaded on Gene Expression Omnibus (link pending).

### Total RNAseq

Total RNA was isolated from cell lines using a phenol:chloroform extraction method. RNA samples were treated with DNase I (Qiagen). Quality control for extract RNA was performed on the Agilent BioAnalyzer (Agilent Technologies, Santa Clara, CA, USA) using the RNA 6000 Nano kit (Agilent Technologies, Santa Clara, CA, USA). Concentration was determined by the Invitrogen Qubit (Thermo Fisher Scientific, Waltham, MA, USA) using the RNA BR reagents (Thermo Fisher Scientific). The TruSeq RNA Sample Prep Kit v2 for Illumina (Illumina, San Diego, CA, USA) was used for cDNA synthesis and adding barcode sequences. The sequencing of the libraries was performed via a paired-end run on a NextSeq500 Illumina sequencer (Illumina). As an average, 25 million reads were collected per sample.

For the RNA-Seq analysis, the sequencing reads were first mapped to the human reference genome (hg38) using the STAR aligner ^80^, ensuring high-quality alignment and accurate mapping. Gene/Isoform expression was then quantified using RSEM, leveraging the Gencode v22 annotation as the gene model to associate reads with genomic features effectively.

To identify differentially expressed genes (DEGs), the limma package was employed, which uses a linear modeling framework to analyze expression data. To account for the multiple comparisons inherent in high-throughput experiments, the Benjamini-Hochberg (BH) procedure was applied to adjust p-values, ensuring control of the false discovery rate (FDR).

### TSS selection

For each putative enhancer RNA (eRNA), the corresponding transcription start site (TSS) was determined based on the publicly available CAGE data produced by the FANTOM consortium. This approach associates eRNAs with experimentally validated TSS. An ad-hoc R script was developed to automate the integration of eRNA coordinates with the CAGE TSS annotations.

### CRISPRi library design

pgRNAs library was designed as previously described in ^81,82^ with a few modifications to better fit CRISPRi (https://github.com/sunandinini/CRISPETa_i-a/tree/bug-fix). 99 positive controls were chosen among validated essential genes^83^ expressed in 43 of our lymphoma cell lines, while 33 neutral controls were selected among genes completely not expressed in any of the 43 lymphoma cell lines.

6-10 independent constructs per each RNA to be tested, 4618 pgRNAs versus eRNAs and lincRNAs and 1207 control pgRNAs, for a total of 5825 constructs. All pgRNA sequences are listed in Table S2. The custom CRISPRi Library was cloned and expanded in outsourcing (Vector Builder Inc.)

### Stable dCas9-ZIM3-expressing cells preparation

VL51 parental or ibrutinib-resistant were infected with the lentiviral vector pHR-UCOE-SFFV-dCas9-mCherry-ZIM3-KRAB (RRID:Addgene_154473). Lentivirus production was carried out by co-transfecting HEK293T cells with 6 mg of dCas9 plasmid, 4.5 mg psPAX2 plasmid, and 1.5 mg of the packaging pVsVg plasmids, using JetPrime (Polyplus). 24 h before the transfection, 3 million HEK293T cells were seeded in a 10 cm dishes. The supernatant containing viral particles was harvested 48, and 72 h after transfection. Viral particles were then concentrated 100-fold by adding 1 volume of Lenti-X Concentrator (Takara) to every three volumes of supernatant. After 12 h at 4 C, the supernatant was centrifuged at 1,500 g for 45 min at 4 C, resuspended in complete medium, and stored at -80 C until use.

For the generation of stable dCas9-expressing cell lines, cells were incubated for 72 h with medium containing concentrated viral preparation and 8 mg/mL polybrene. MCherry+ cells were sorted several times before obtaining cell lines able to sustain dCas9-ZIM3-KRAB expression and cultured in the presence of additional nonessential amino acids and higher content of fetal bovine serum. Cells were also tested for dCas9 expression by western blotting, and the protein’s functionality was proven by infection with sgRNA directed against MALAT1 TSS (pDECKO_Malat1_C, RRID: Addgene_72622). Two independent dCas9 clones for parental and two for resistant cells were selected as biological replicates to infect in parallel during the screen.

### Lentiviral library titration

The functional titer of the CRISPRi library was evaluated, infecting a small scale of dCas9-ZIM3-KRAB-mCherry cells with an increasing amount of virus. After 48 and 72 hours, cells were checked for GFP expression by FACS, and the volume of virus per million cells was calculated based on the infection that achieved 30% GFP+ cells.

### elncRNA CRISPRi screen

Fifty million cells were infected with the CRISPRi Library at an MOI of 0.3 and then selected by puromycin exposure for two days, followed by a two-day washout period. One-third of the cells, corresponding to a minimum of 3 million, were saved as T0 to obtain the baseline representation of the pgRNAs distribution. Then, 3 million mutant cells per experiment group were selected by exposure to ibrutinib or DMSO and culture under drug pressure for 14 days. The number of cells was calculated to represent 500 cells per each of the 6000 pgRNAs. The dose chosen for the treatment was 500 nM of ibrutinib (a sub-active dose equivalent to 1/10 of the IC50) in parental cells. In parallel, mutant cells derived from the resistant VL51 were treated with 5 µM of ibrutinib (equivalent to the IC50 of the parental and an inactive dose in these cells). In parallel, the same number of cells were exposed to DMSO to highlight molecules relevant to cell survival independent of the drug response.

To achieve 100x coverage of the library, at least 5 µg of gDNA per sample was amplified using appropriate primers flanking the construct region containing the sgRNA. PCR products were purified and then sequenced by amplicon-based next-generation sequencing, 75 bp single-end, using NextSeq500 sequencer. The sgRNAs were mapped, and their distribution was analyzed in all the experimental groups and in the baseline controls. MAGeCKFlute software ^84^ was applied to perform a quality check of the CRISPRi screen.

### Lentivirus and cell line production for competition assay

pgRNAs targeting TSS of elncRNAs selected for further validation were cloned in the pDECKO mCherry vector (RRID:Addgene_78534) as previously described ^82^. All the pgRNA guides for CRISPRi validations are reported in Table S12. Lentiviral supernatant for each pgRNA construct was produced as described above. After 72h infection with pgRNA-pDECKO lentiviral supernatants, parental or ibrutinib-resistant VL51 or Jeko1 cells were selected with puromycin (1.5 mg/mL) for two days.

Cells stably expressing pgRNAs and mCherry were then infected with dCas9-Zim3-GFP (RRID: Addgene_188778). After 72 h, cells were analyzed by FACS to determine the percentage of mCherry-GFP double-positive cells at T0. Then, the cells were split into equal amounts and exposed to DMSO or ibrutinib, and tested every three days to measure the survival of double-positive cells compared to cells expressing only pgRNAs. After 14 days double-positive cells were sorted to obtain pure polyclonal populations, to test for ibrutinib sensitivity and for transcriptomic analysis.

The polyclonal populations were also seeded at limiting dilutions to obtain monoclonal cell lines, which were screened for representativeness of the polyclonal cell lines and used for additional validation experiments.

### A-to-I editing analysis from tot RNAseq

To analyze A-to-I RNA-editing events, we utilized the RED-ML algorithm ^49^, a method specifically designed for RNA-editing detection. The analysis began with the aligned sequencing data in BAM format, ensuring that high-quality mapped reads were used as input. RED-ML systematically identified putative RNA-editing sites by distinguishing true A-to-I editing events from sequencing artifacts and single nucleotide polymorphisms (SNPs), leveraging its model for increased accuracy and specificity. Following the identification of RNA-editing events, we annotated the edited sites with gene annotations to associate each editing event with its corresponding gene. Additionally, we classified the events based on their genomic context, determining whether they occurred within gene bodies (3’UTR, 5’UTR, exonic or intronic regions) or in promoter/downstream/intergenic regions. To further assess the functional implications of these RNA-editing events, particularly in protein-coding genes, we utilized ANNOVAR to evaluate their potential impact on coding sequences. Also, editing sites were annotated based on genomic location, and if they belonged to an Alu sequence or not. Editing sites were considered sequence-specific if the same nucleotide was edited in at least three samples. Differential RNA editing analysis was performed using the voom/limma R package ^85^. Sequence-specific edited sites were considered for further analyses. Differentially edited sites were defined as those with an empirical Bayes corrected (Benjamini-Hockberg procedure) p-value <0.05.

### Exon Intron Splicing Analysis

Post-transcriptional modulation of transcripts was estimated as previously described ^86^

### RNA fractionation

RNA fractionation was performed as previously described ^18^. Briefly, 5 million cells were lysed with RSB-100 buffer containing digitonin to permeabilize cells and recover cytoplasmic RNA. Nuclei were lysed by RSB-100 buffer after the addition of 0.5% Triton-X to collect nuclear RNA by ultracentrifugation. The insoluble fraction was resuspended in nuclear lysis buffer and sonicated to extract chromatin-associated RNA.

### DRS

To identify full-length spliced elncRNAs and alternative protein-coding transcripts at a genome-wide level, all nuclear transcripts, polyadenylated or not, were enriched. 3 µg of RNA extracted from two monoclonal ADARreg_kd and two negative control KLK12_kd ibrutinib-resistant cell lines were used for ribodepletion by NEBNext rRNA Depletion Kit. Then, synthetic poly-A tails were added to non-polyadenylated transcripts by NEB E. coli Poly(A) Polymerase reagent, for library preparation compatibility. For direct sequencing of native RNA transcripts and modifications, libraries were prepared with the PCR-free Direct RNA Sequencing (DRS) Kit (SQK-RNA004, Oxford Nanopore) starting from 300 ng of RNA, purified after the previous enzymatic steps and run by Promethion 24 (Oxford Nanopore), two biological replicates per condition. Around 30 million reads per sample were collected, running a single RNA sample per flow cell. Similarly, cytosolic RNA was used for library preparation but starting from 1 µg of unprocessed cytosolic fractions.

Oxford Nanopore Technologies (ONT) long-read sequencing data were processed using Dorado (https://github.com/nanoporetech/dorado). In particular, Dorado was used for basecalling and for generating POD5 files directly from raw signals, followed by read alignment to the human reference genome (GRCh38/hg38) using the integrated alignment module, which internally utilizes minimap2 (https://github.com/lh3/minimap2). Dorado’s model-based detection framework was also applied to call RNA modifications, specifically N6-methyladenosine (m^6^A) and adenosine-to-inosine (A-to-I) edits using the rna004_130bps_hac@v5.2.0 model. To assign modification events to transcript isoforms, we developed an in-house script that links modification calls to isoform-resolved alignments. Gene-and isoform-level quantification, as well as de novo transcript discovery, were performed using Bambu ^87^ , which provides an integrated approach for long-read–based transcript reconstruction and abundance estimation.

### Quantitative Reverse Transcriptase Polymerase Chain Reaction (qRT-PCR)

Strand-specific quantitative RT-PCR (qRT-PCR) was performed using SuperScript III Platinum SYBR Green One-Step qRT-PCR Kit (Invitrogen). Data were analyzed according to the manufacturer’s instructions and the comparative CT method (ΔΔCT method) then normalized to GAPDH or β-actin as reference genes. Statistical significance was determined using a two-tailed t-test with a threshold of p< 0.05. Primer sequences are reported in Supplementary Table S12.

### Immunoblotting

Protein extraction, separation, and immunoblotting were performed as previously described. Nuclear/cytosol fractionation was performed by Nuclear/Cytosol fractionation kit (BioVision, Cat# K266-100) according manufacturer instructions. The following antibodies were used: anti-ADAR1 (D7E2M) (Cell Signaling Technology Cat# 14175, RRID:AB_2722520), Anti ADARB1 (Sigma-Aldrich Cat# SAB1405426, RRID:AB_10740297), anti-IRF3 (D-3) (Santa Cruz Biotechnology Cat# sc-376455, RRID:AB_11151578) anti-NF-κB p65 (C22B4) (Cell Signaling Technology Cat# 4764, RRID:AB_823578), anti-phospho-NFkB p65 (Ser536) (Cell Signaling Technology Cat# 3033, RRID:AB_331284), anti-vimentin, anti-GAPDH (FF26A) from eBioscience, secondary mouse (NA931V) and rabbit (NA934V) antibodies from GE healthcare. The data were analyzed using Fusion Solo software.

### Immunoprecipitation

Nuclei from 10 million cells were isolated using RSB-100 buffer containing digitonin and a phosphatase and protease inhibitor cocktail. They were then washed and lysed in 1 mL of 50 mM Tris (pH 7.5), 150 mM NaCl, 1% Triton X-100, 1 mM EDTA, and 1 mM EGTA. The insoluble fraction was removed by centrifugation, 25 μL of the supernatant saved as input, and the rest split in two for the overnight incubation with 5 μg antibody RPS3 (Proteintech-66046-1-IgG) or 5 ug μg of mouse IgG2a isotype control (Proteintech 66360-2-IgG-150UL). The day after protein complexes were recovered by magnetic protein A beads (20 μL, 2h 4°C in rotation), then washed three times with lysis buffer and then protein complexes were eluted in 20 μL of Laemmli buffer, boiled 10’ at 95°C and separated as previously described. For the blotting anti-phospho-NFkB p65 (Ser536) (Cell Signaling Technology Cat# 3033, RRID:AB_331284) and RPS3 (Proteintech-66046-1-IgG) were used and detected by Clean-Blot IP Detection Reagent (HRP) (Thermo Scientific™, cat #21230) to avoid the cross reactivity of the heavy and light chains of the bait antibody.

### Immunofluorescence and confocal microscopy

Ibrutinib resistant VL51 cells (100,000 cells/well) were seeded on poly-L-ornithine coated 18-well IBIDI chambers (81816, IBIDI GmbH), then fixed for 20 min with Paraformaldehyde (PFA) 4% at room temperature (RT). Cells were permeabilized with PBS + 0.1% Triton X-100 for 10 min at RT. To block unspecific staining, samples were blocked for 1 h with 5% Bovine Serum Albumin (BSA) at RT before staining. Antibodies were diluted in 5% BSA. Samples were incubated overnight at 4°C with primary antibody anti IRF3 (D-3) (Santa Cruz Biotechnology Cat# sc-376455, RRID:AB_11151578), anti-ADARB1 (Sigma-Aldrich Cat# SAB1405426, RRID:AB_10740297), anti-ADAR1 (Santa Cruz Biotechnology Cat# sc-73408, RRID:AB_2222767), or anti-NF-κB p65 (C22B4) (Cell Signaling Technology Cat# 4764, RRID:AB_823578). Secondary goat antibody anti-mouse IgG labelled with Alexa647 (A-21235, Thermo Fisher Scientific Cat# A-21235, RRID:AB_2535804) 1 h at RT in the dark. Slides were counterstained after 3 washes of PBS with 0.3Wμg/mL 4,6-diamidino-2-phenylindole (Sigma-Aldrich). Images were acquired on a Leica SP5 with an objective with ×63 magnification. Protein quantification and nuclear/ cytosol ratio were evaluated by ImageJ (RRID:SCR_003070) software. At least three images per condition, including approximately 30-50 cells each, were used for each quantification.

### Co-culture with NK cells

PBMCs were collected from healthy donors through density gradient separation (Ficoll Paque Plus, Merck). NK isolation was performed using the Easy-Sep Human NK Cell Isolation kit (STEMCELL Technologies) following manufactural instructions. Freshly isolated NK cells were cultured with RPMI 1640 medium supplemented with IL-2 (2.5ng/ml; PeproTech) and IL-15 (10ng/ml; PeproTech).

### Single-cell RNAseq data

The single-cell sequencing dataset (GSE111014) was obtained from GEO and is associated with the referenced publication ^51^. The re-analysis of the dataset files (GSE111014_barcodes.tsv.gz, GSE111014_genes.tsv.gz, and GSE111014_matrix.mtx.gz) was conducted from scratch using the Seurat package in R. The raw data were pre-processed to generate a Seurat object, followed by quality control, normalization, and feature selection steps to identify variable genes. Dimensionality reduction was performed using principal component analysis (PCA) and visualized through uniform manifold approximation and projection (UMAP). For cell-type annotation, the SingleR algorithm was applied, leveraging reference datasets to predict cell identities. The output of SingleR was then subjected to a manual revision to refine and ensure the accuracy of the annotations, combining algorithmic insights with domain expertise. B cells were subdivided in ADARreg ^high^-like or ADARreg ^low^-like based on the signature score calculated by R package AUCell using IBRr VL51 ADARreg knockdown differentially expressed genes. NK cell subsets were defined based on cytotoxicity, inflammatory and stress-related gene sets as previously described^52^. Cell-cell interactions at pretreatment time points in patient CLL8 and CLL6 were analyzed by CellChat.

## Data mining

Functional analysis was performed using GSEA^88^ (Gene Set Enrichment Analysis) on genes ranked by t-statistic as calculated by Limma test from differential gene expression analysis. Gene sets were considered significantly enriched if FDR<0.05.

Gene ontology analysis was performed using the g-Profiler^89^ webtool. The p-value for pathway enrichment was computed using a Fisher’s exact test multiple-test correction was applied.

Functional impact of differential isoform expression was performed by tappAS^32^ software using a functional iso-transcriptomic approach to retrieve functional annotation features at both RNA and protein levels.

Semantic clustering analysis was performed on long lists of gene ontology terms using Revigo^90^. Statistical methods were implemented in R and executed within the Rscript environment.

## Supplemental information

Figure S1 (related to Figure 2): Functional screening of elncRNAs linked to BCR blockade

Figure S2 (related to Figure 2): lncPCED1b and ADARreg elncRNAs genomic loci Figure S3 (related to Figure 3): lncPCED1b and ADARreg knockdown biological effects Figure S4 (related to Figure 4): A-to-I RNA editing evaluated by total RNAseq

Figure S5 (related to Figure 5): Transcriptional and post-transcriptional impact of lncPCED1b and ADARreg silencing evaluated by total RNAseq.

Figure S6 (related to Figure 6): Transcriptional and post-transcriptional impact of ADARreg silencing evaluated by DRS on subcellular compartments.

Figure S7 (related to Figure 7): Modulation of key RNA isoforms of the immune-suppressive transcriptional program of lymphomas by ADARreg.

Figure S8 (related to Figure 7): Crosstalk between NK cell and tumor correlated with ADARreg signature in CLL patients.

Figure S9 (related to Figure 7): proposed model

**Table S1 (separate file, related to Fig.1): VL51_Ibru vs parental VL51 specific enhancers and DEGs**

Ibrutinib-resistant versus parental VL51 specific enhancers detected by ChIPseq and annotated with differentially expressed genes, resulting from the comparison of gene expression profiles of VL51-Ibru versus parental VL51 and falling within a 40kb window from the peak. LogFC represents modulation for each gene in ibrutinib-resistant vs parental.

**Table S2 (separate file, related to Fig.2): Composition of elncRNA SMZL CRISPR-i library**

Composition of elncRNA SMZL CRISPR-i library. Sequences of both sgRNAs of the pair binding each target with relative chromosomal position are listed.

**Table S3 (separate file, related to Fig.2):** CRISPRi screen

CRISPRi screen enrichment scores (beta) of each sample respect the time 0. Targets are considered statistically significant enriched for fdr<0.2 (green)

**Table S4**: **(separate file, related to Fig.4): A-I-RNA editing**

Limma test performed on A-to-I RNA editing for each edited site detected by RED-ML. LogFC represents modulation for each editing site in ibrutinib-resistant vs parental cells (RESacq) or in parental (PAR) or ibrutinib-resistant (RES) cells knockdown for elncRNA of interest vs the negative control.

**Table S5 (separate file, related to Fig.5): DEG ADARreg_lncPCED1b_vs_KLKL12kd**

Limma test performed on gene expression profiles of ibrutinib-resistant vs parental cells (IBRr_KLK12kd vs PAR_KLK12kd) or in ibrutinib-resistant cells knockdown for ADARreg or lncPCED1b vs the negative control. LogFC represents modulation for each significantly differentially expressed gene (adj.P<0.05) in the indicated comparison

**Table S6 (separate file, related to Fig.5): GSEA**

Genesets significantly enriched in at least one of the following comparisons: ibrutinib-resistant versus parental cells, ibrutinib-resistant ADARreg or lncPCED1b knockdown versus negative control. Enrichment score (NES) and fdr calculated by GSEA software are indicated for each geneset and each comparison.

**Table S7 (separate file, related to Fig.5): Post-transcriptionally modulated transcripts_EISA**

D exons and D introns measured by EISA in the comparison of ibrutinib-resistant cells with ADARreg or lncPCED1b knockdown versus negative control. Transcripts are considered post-transcriptionally modulated if Dex/Dint in ADARreg or lncPCED1b vs KLK12_kd have p<0.05 in Limma test.

**Table S8 (separate file, related to Fig.S6): editing of alternative transcripts**

List of genes with transcriptional isoforms modulated in opposite direction (opposite LogFC and adj p-value <0.05 in Limma test performed on quantification per isoforms of total RNAseq in ibrutinib-resistant ADARreg or lncPCED1b knockdown versus control cells). Additional information about significantly differential intronic or 3’ UTR editing events is provided.

**Table S9 (separate file, related to Fig.6): DIU DEA summary**

TappAS summary of differential isoform usage (DIU) and differential expression analysis (DEA) in nuclear transcripts quantified by full-length reads in DRS.

**Table S10 (related to STAR Methods): oligos for CRISPRi sequencing and validation experiments**

Sequences of pgRNAs cloned in lentiviral vector for stable silencing of elncRNAs, antisense oligonucleotides used for transient silencing, primers to prepare CRISPRi library amplicons for sequencing, and primers to detect transcript expression by qRT.

## Notes

### Summary of Updates

Direct RNA sequencing was performed on subcellular fractions. This approach enabled a precise evaluation of the effects of ADARreg knockdown on RNA modification landscapes and on the expression and processing of transcriptional isoforms.

