## Supplementary material for "Enhancer lncRNA LOC730338 modulates BCR signaling and immune evasion in lymphoma by regulating RNA homeostasis": Supplemetal Figures

**Figure S1 (related to Figure 2):** Functional screening of elncRNAs linked to BCR blockade

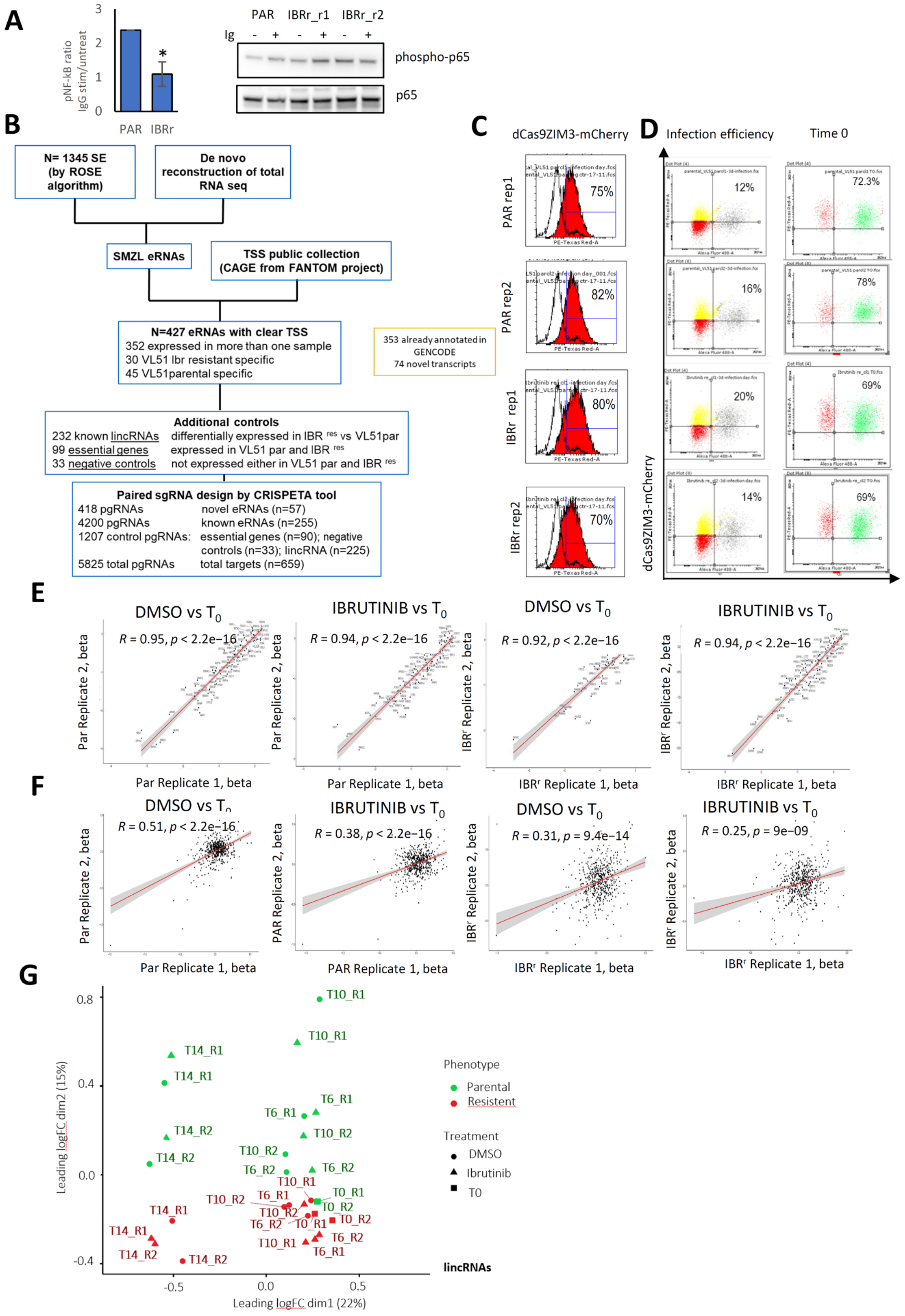

**(A)** Quantification (left) of phosphorylation of NF-kB detected by western blotting (right) in VL51 or VL51-Ibru cells exposed to IgG with respect to unstimulated cells.

**(B)** Pipeline for the selection of e-lncRNAs and controls to include in the CRISPRi library.

**(C)** dCas9 ZIM3-GFP expression measured by flow cytometry in two replicates of VL51 or VL51-Ibru (red) respect not infected control.

**(D)** Efficiency of infection and then selection of dCas9-ZIM3-GFP+ cells with pgRNA lentiviral library quantified by mCherry fluorescence by FACS before, left, and after, right, puromycin selection.

**(E)** Scatter plot representing the b values of pgRNAs targeting the ncRNAs in the two replicates per each experimental condition. Pearson correlation test verifies the concordance of replicates.

**(F)** Scatter plot representing the b values of pgRNAs targeting the positive controls in the two replicates per each experimental condition. Pearson correlation test verifies the concordance of replicates

**(G)** MDS plots showing the distribution of samples at the beginning of CRISPRi screen (T0, squared labels) or after 6 (T6), 10 (T10) or 14 (T14) days, based on the drop out of lincRNAs. Parental cells are represented in green, ibrutinib-resistant in red. Samples exposed to the drug are labelled with triangle, to the vehicle with the circle.

**Figure S2 (related to Figure 2):** lncPCED1b and ADARreg elncRNAs genomic loci

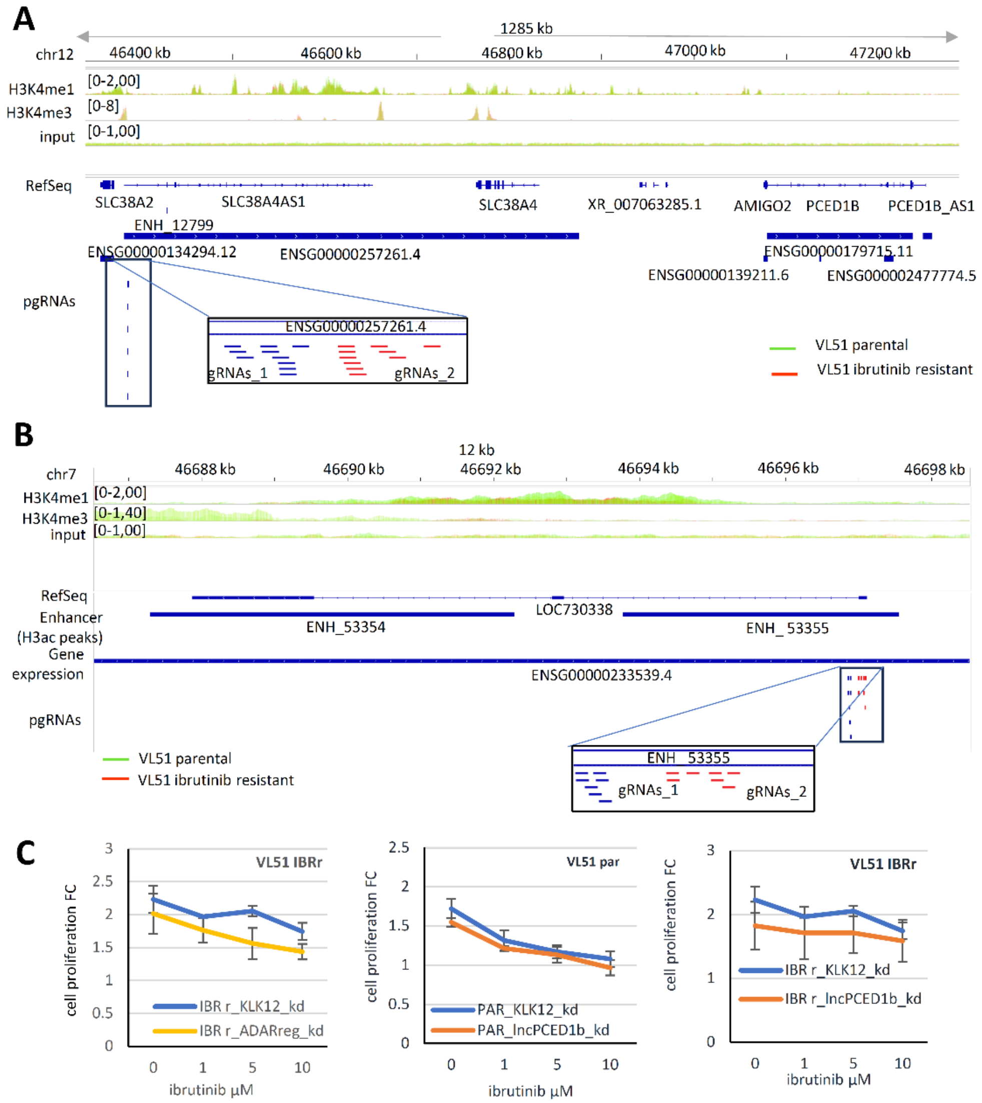


- **(A)** Schematic representation of RP11-96H19.1 locus. H3K4me1, H3K4me3, and input tracks are overlays of VL51 (green), and VL51-Ibru (red) replicates. Genomic positions of acetylated H3 peaks and expressed transcripts in ibrutinib-resistant or parental cells are indicated. Bottom, close-up of pgRNAs targeting RP11-96H19.1 TSS.
- **(B)** Schematic representation of AC011294.3 locus. H3K4me1, H3K4me3, and input tracks are overlays of parental (green), and ibrutinib-resistant derivatives (red) replicates. Genomic positions of acetylated H3 peaks and expressed transcripts in ibrutinib-resistant or parental cells are indicated. Bottom, close-up of pgRNAs targeting AC011294.3 TSS.
- **(C)** MTT assay performed at 72h of ibrutinib treatment in GFP+ mCherry+ sorted cells, 2 weeks after dCas9 lentiviral infection in ibrutinib-resistant VL51 cells stably expressing mCherry and pgRNA targeting the negative control KLK12, or the elncRNA, ADARreg, top, or lncPCED1b in parental, middle, or in ibrutinib-resistant cells, bottom.

**Figure S3 (related to Figure 3):** lncPCED1b and ADARreg knockdown biological effects

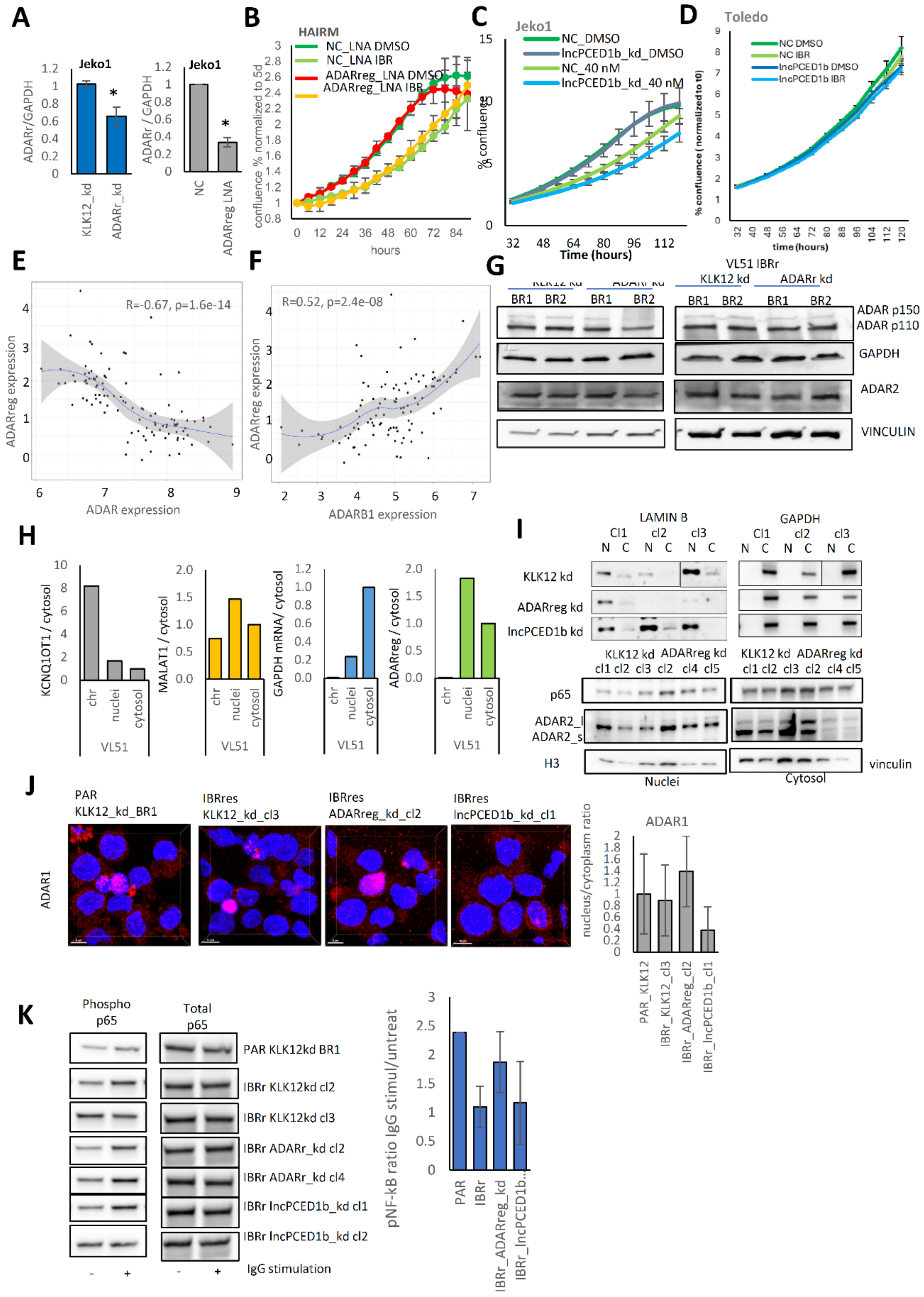

- **(A)** ADARreg expression normalized to GAPDH in Jeko1 stable silenced for KLK12 negative control or ADARreg (left), or 4 days after nucleofection of Jeko1 with ADARreg LNA or negative control (right).
- **(B)** Growth curve of HAIRM seeded 4 days after ADARreg LNA or negative control electroporation and built measuring cell confluence over 4 days under exposure to ibrutinib or vehicle
- **(C)** Growth curve of Jeko1 seeded 4 days after lncPCED1b LNA or negative control electroporation and built counting cells over 5 days under exposure to ibrutinib or vehicle.
- **(D)** Growth curve of Toledo seeded 4 days after lncPCED1b LNA or negative control electroporation and built counting cells over 5 days under exposure to ibrutinib or vehicle.
- **(E)** Pearson correlation of ADARreg and ADAR in 102 MCL patients.
- **(F)** Pearson correlation of ADARreg and ADARB1, in 102 MCL patients.
- **(G)** Western blotting of ADAR1 and ADAR2 proteins in two independent biological replicates of Jeko1, left, or ibrutinib-resistant VL51, right, KLK12 kd or ADARreg kd. GAPDH and vinculin are housekeeping controls for ADAR1 and ADAR2.
- **(H)** Subcellular localization of ADARreg. KCNQ1OT1, MALAT1 and GAPDH mRNA were used as positive control of chromatin-associated, nuclear, or cytoplasmic RNA, respectively. A representative experiment is shown.
- **(I)** Nuclear and cytoplasmic protein distribution evaluated by western blotting after subcellular fractionation of three independent monoclonal ibrutinib-resistant ADARreg_kd or KLK12_kd VL51 cells. Top, fractionation quality check by lamin B and GAPDH as respectively nuclear and cytosolic proteins. Bottom, p 65 and ADAR2 subcellular distribution, H3 and vinculin were used for loading controls.
- **(J)** Left, Immunofluorescence of ADAR1 in ibrutinib-resistant KLKL12 kd (negative control) or ADARreg kd monoclonal cell lines. A polyclonal parental KLK12kd VL51 cell line was compared as an additional baseline control. DAPI stained nuclei, blue, Alexa647-conjugated antibody stained ADAR1, red. A blue and red fluorescence overlay was reported. Two examples at single-cell magnification were included. Right, graph with nuclei/ cytoplasm ratio of ADAR2 integrated density.
- **(K)** Phosphorylation of NF-kB detected by western blotting (left) and quantified (right) in parental or ibrutinib-resistant cells after KLK12, ADARreg, or lncPCED1b kd, exposed to IgG respect to unstimulated cells.

**Figure S4 (related to Figure 4):** A-to-I RNA editing evaluated by total RNAseq

**A**

| source | term_name | adj_p_value | negative_log10_ of_adj_p_value | term_size | query_size | intersections |
|---|---|---|---|---|---|---|
| KEGG | Antigen processing and presentation | 9.53E-05 | 4.020739024 | 69 | 77 | CTSS,CTSB,HLA-B,HLA-DPB1 |
| KEGG | Epithelial cell signaling in Helicobacter pylori infection | 0.000172 | 3.763250855 | 70 | 88 | F11R,LYN,PLCG2,JAM3 |
| KEGG | Epstein-Barr virus infection | 0.000272 | 3.566184496 | 198 | 77 | LYN,HLA-B,IFNAR2,PLCG2,HLA-DPB1 |
| KEGG | Pathways in cancer | 0.001569 | 2.80438781 | 527 | 70 | TFG,TRAF1,PGF,LAMC1,IFNAR2,PLCG2 |
| REAC | DNA Repair | 9.11E-06 | 5.040453817 | 329 | 91 | ERCC6,FAAP24,UBE2L6,POLH,RFC4,CLSPN,RMI2,BRCA1 |
| REAC | Cell Cycle | 1.73E-05 | 4.761827146 | 677 | 91 | HMMR,TUBGCP6,LYN,DYNC1H1,OIP5,RFC4,SFI1,CLSPN,RMI2,BRCA1 |
| REAC | Adaptive Immune System | 1.78E-05 | 4.748479725 | 809 | 77 | CLEC2D,CTSS,HMGB1,CTSB,UBE2L6,LYN,DYNC1H1,HLA-B,PLCG2,HLA-DPB1 |
| REAC | MHC class II antigen presentation | 0.005212 | 2.282997409 | 123 | 77 | CTSS,CTSB,DYNC1H1,HLA-DPB1 |

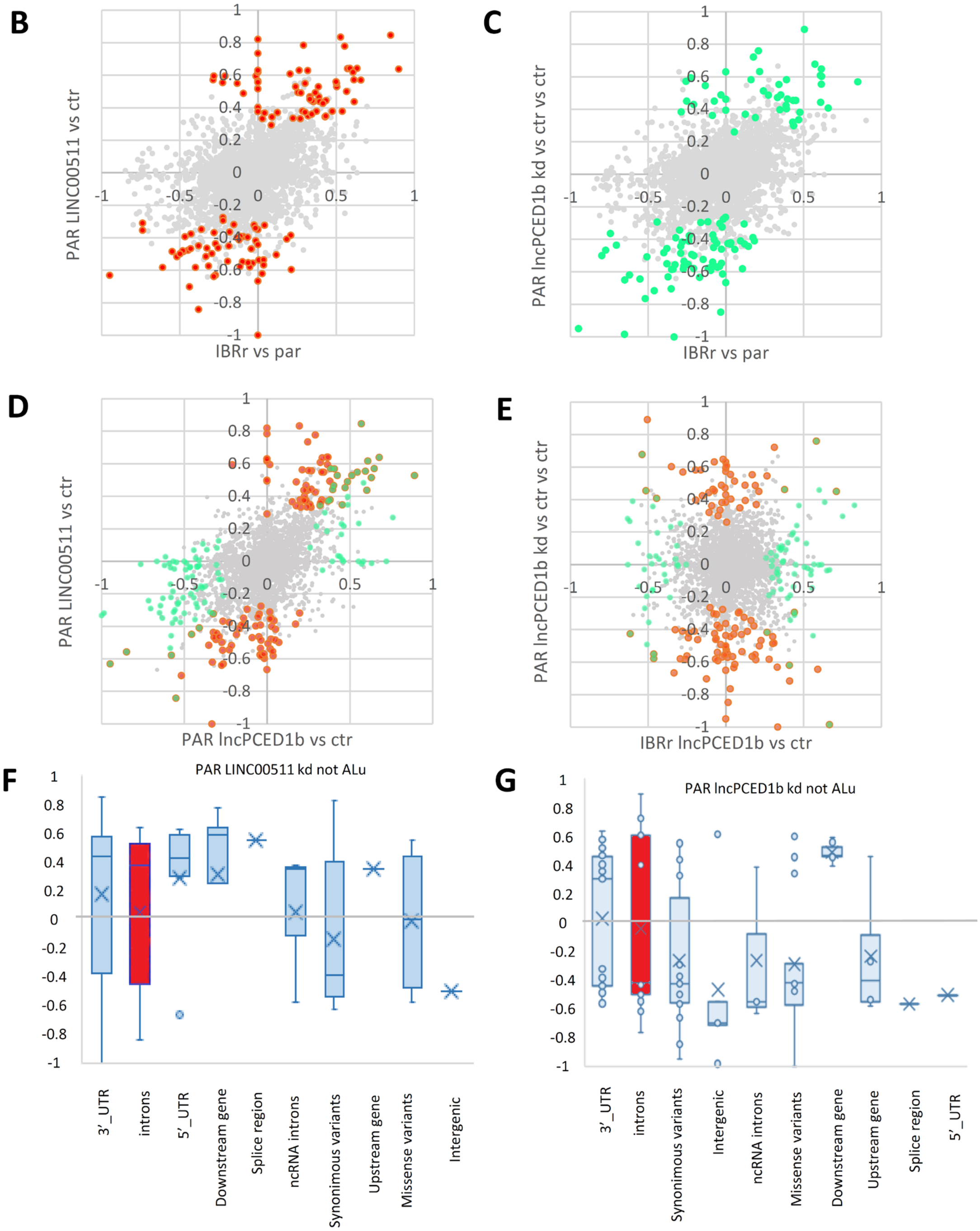

**(A)** Gene ontology analysis of genes differentially edited after knockdown both of ADARreg and lncPCED1b in ibrutinib-resistant cells

**(B)** Scatter plots of editing site Log FC in the comparison resistant versus parental and parental VL51, LINC00511kd cells versus KLK12kd control. Red dots represent editing sites statistically significant in the comparison LINC00511 kd versus control.

**(C)** Scatter plots of editing site Log FC in the comparison resistant versus parental and parental VL51, lncPCED1b kd cells versus KLK12kd control. Green dots represent editing sites statistically significant in the comparison lncPCED1b kd versus control.

**(D)** Scatter plots of editing site Log FC in the comparison parental VL51, lncPCED1b kd cells versus KLK12kd control and LINC00511 kd versus control. Orange dots represent editing sites statistically significant in the comparison LINC00511 kd versus control, light green dots in lncPCED1b kd versus control, and dark green in both.

**(E)** Scatter plots of editing site Log FC in the comparison ibrutinib resistant, VL51, lncPCED1b kd cells versus KLK12kd control and parental VL51, lncPCED1b kd cells versus KLK12kd control. Orange dots represent editing sites statistically significant in the comparison of lncPCED1b kd versus control in parental cells, light green dots in ibrutinib-resistant cells, and dark green in both.

**(F)** Bar plots showing the log fold change distribution of differentially not-Alu edited sites within different genomic locations, in the comparison of parental VL51 cells LINC00511_kd respect to the KLK12kd control cells (pvalue< 0.05, Limma test, median value).

**(G)** Bar plots showing the log fold change distribution of differentially not-Alu edited sites within different genomic locations, in the comparison of parental VL51 cells lncPCED1b_kd respect to the KLK12kd control cells (pvalue< 0.05, Limma test, median value)

**Figure S5 (related to Figure 5):** Transcriptional and post-transcriptional impact of lncPCED1b and ADARreg silencing evaluated by total RNAseq.

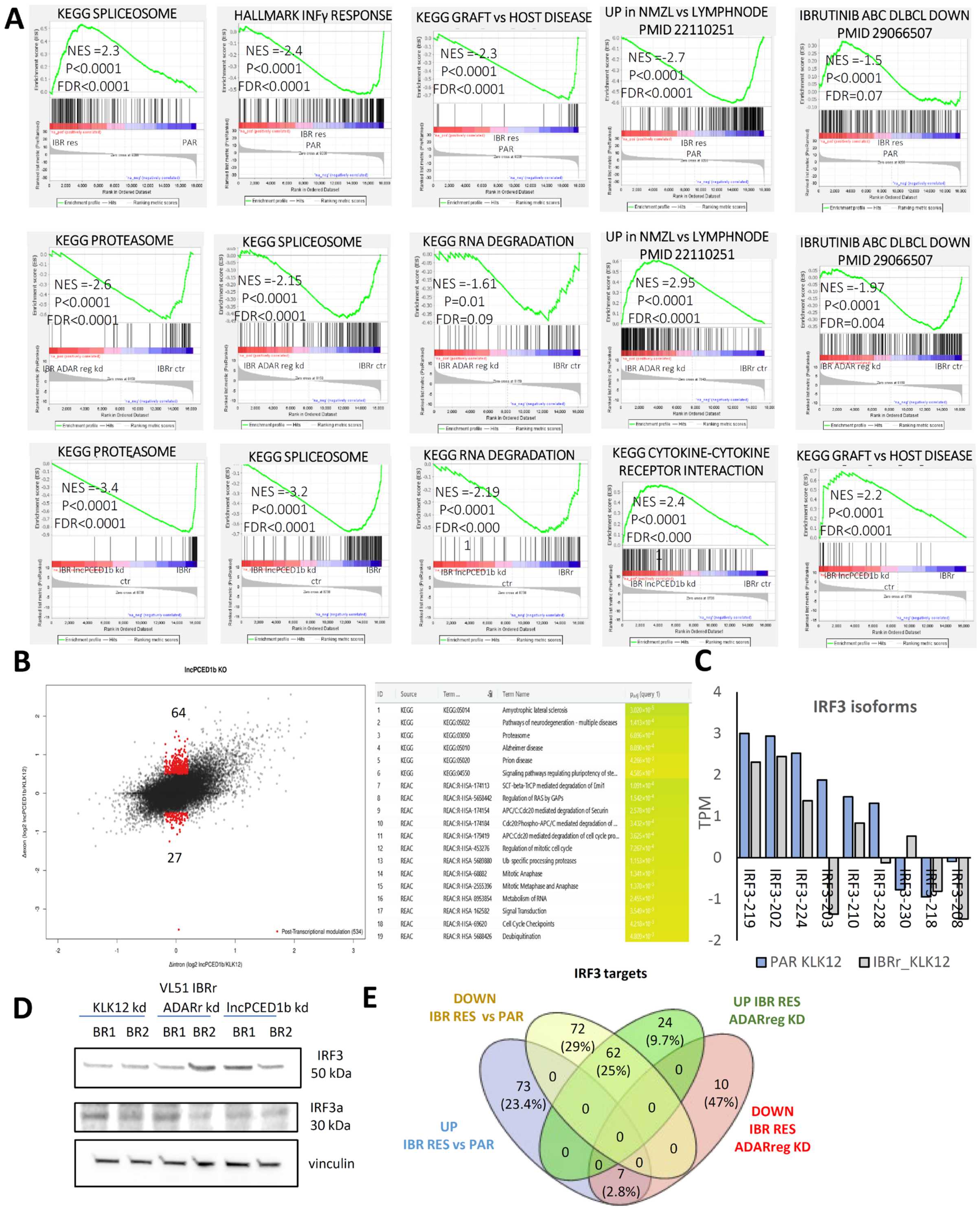

- **(A)** Preranked GSEA of RNAseq data in ibrutinib-resistant VL51 KLK12 kd respect to parental VL51 KLK12 kd, top, after ADARreg knock down in ibrutinib-resistant VL51, middle, or lncPCED1b knock down, bottom.
- **(B)** Top, Scatter plots representing the comparison of changes in intronic (Δintron) and exonic (Δexon) reads per each expressed transcript in ibrutinib-resistant VL51 lncPCED1b kd respect to KLK12 kd negative control. Δexon/ Δintron=1 reflects transcriptional modulation, Δexon/ Δintron≠1 reflects post-transcriptional modulation. Red dots represent robust post-transcriptionally modulated transcripts. P value<0.05. Bottom, gProfiler gene ontology analysis of transcripts significantly modulated at post-transcriptional level after lncPCED1b kd.
- **(C)** All IRF3 isoforms measured in total RNAseq performed in parental and ibrutinib-resistant VL51 negative control cells
- **(D)** Western blotting of IRF3 isoforms, IRF3 and IRF3a, in two independent biological replicates of ibrutinib-resistant VL51 KLK12 kd or ADARreg kd. Vinculin is a housekeeping control.
- **(E)** Venn diagram of IRF3 target genes significantly modulated in the comparison ibrutinib resistant versus parental VL51 cells or after ADARreg kd in ibrutinib resistant cells.

**Figure S6 (related to Figure 6):** Transcriptional and post-transcriptional impact of ADARreg silencing evaluated by DRS on subcellular compartments.

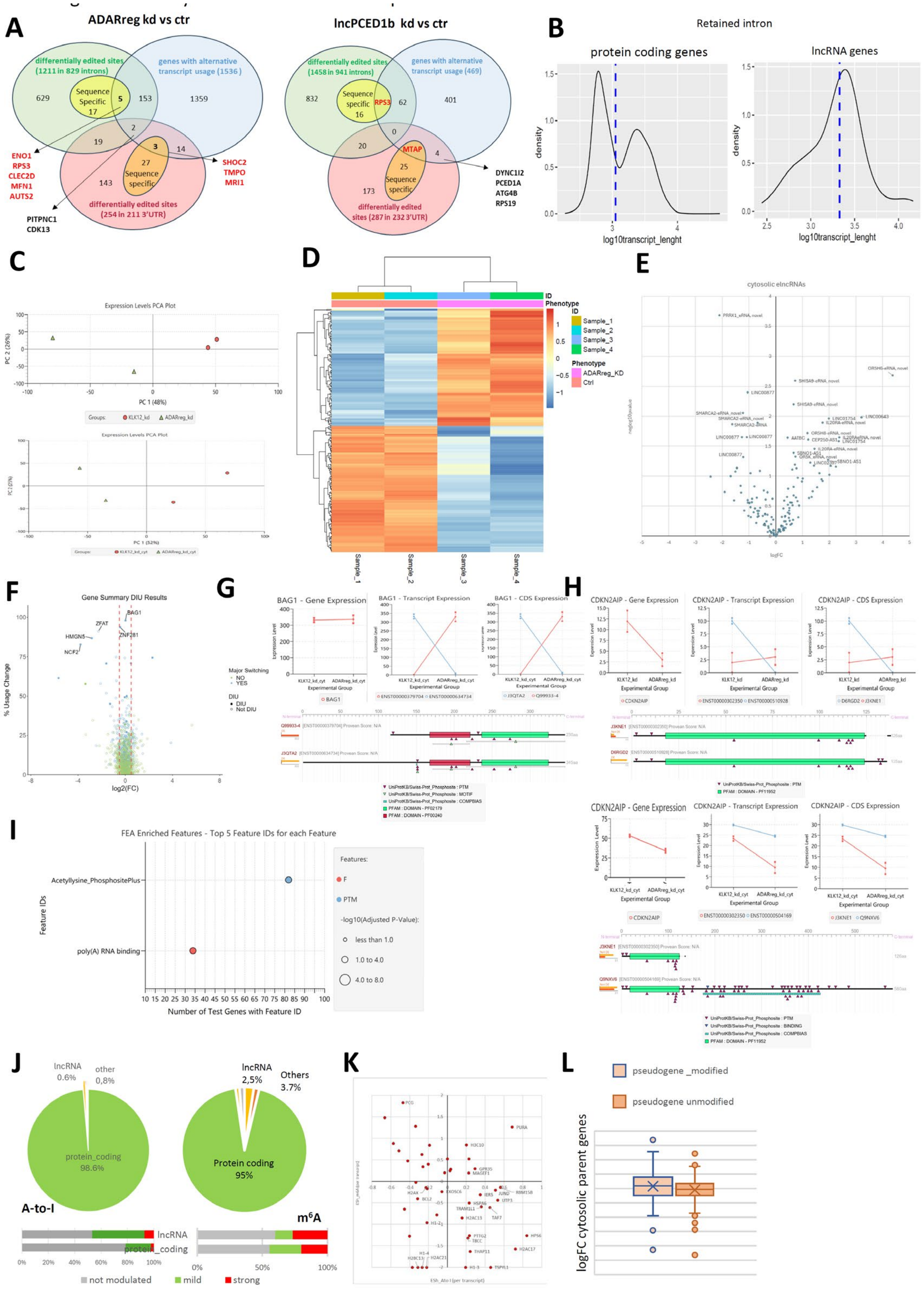

**(A)** Venn diagram intersecting total and sequence-specific edited transcripts significantly modulated in introns or 3'UTR and alternative transcriptional isoforms in ibrutinib-resistant VL51 after ADARreg kd, left, or lncPCED1b kd.

**(B)** Length distribution of nuclear retained intron transcripts transcribed from protein coding, left, or lncRNA genes, right.

**(C)** MDS plots show sample clustering in the DRSeq full-length read quantification in the nucleus, top, and in the cytosol, bottom.

**(D)** Heat map on nuclear top-modulated transcripts, quantified by full-length reads, in the comparison of two biological replicates per condition, ADARreg _kd versus control KLK12_kd ibrutinib-resistant cells.

**(E)** Volcano plot of elncRNAs in the cytosol. Labels indicate the gene symbol of the most significant.

**(F)** Total usage change (i.e., expression redistribution between isoforms) vs log-transformed values of gene expression fold change between ADARreg kd and control in the cytosol. Genes with a major isoform switch are represented in blue. Labels indicate genes with the highest total usage change. Full dots represent genes with significant differential isoform usage (DIU).

**(G)** From left to right: gene, transcript, and protein-level expression charts for the BAG1 gene in the two conditions in the cytosol (ADARreg kd and control KLK12 kd, each containing two replicates, squares indicate replicate expression, points indicate mean replicate expression), and tappAS graphical representation of protein-level annotations. Colored boxes represent protein domains.

**(H)** From left to right: gene, transcript, and protein-level expression charts for the CDKN2AIP gene in the two conditions (ADARreg kd and control KLK12 kd, each containing two replicates, squares indicate replicate expression, points indicate mean replicate expression), in the nucleus, top, and in the cytosol, bottom. TappAS graphical representation of protein-level annotations. represent protein domains by colored boxes.

**(I)** Functional enrichment of DIU genes for GO terms (Fisher's exact test, multiple testing correction by Benjamini-Hochberg method, FDR < 0.05). Dot color indicates the GO term aspect, while dot size indicates significance level.

**(J)** Top, distribution of A-to-I and m6A sites, detected in cytosolic full-length read transcripts by DRseq and assigned to different biotypes. Bottom, percentage of A-to-I and m6A sites significantly different in the comparison ADARreg kd versus control using Cohen's index (Esh) to quantify effect size (not modulated ($|Esh|<0.2$, mild ($0.2<|Esh|<0.5$), strong ($|Esh|>0.5$))

**(K)** Scatter plot of transcript-aggregated Cohen indexes, Esh, for A-to-I and m6A of nuclear protein coding transcripts.

**(L)** LogFC distribution of cytosolic transcripts derived from parent genes, whose corresponding nuclear pseudogenes were differentially modulated in terms of both $m^6A$ and A-to-I (orange) or $m^6A$ only (blue) after ADARreg kd.

**Figure S7 (related to Figure 7):** Modulation of key RNA isoforms of the immune-suppressive transcriptional program of lymphomas by ADARreg.

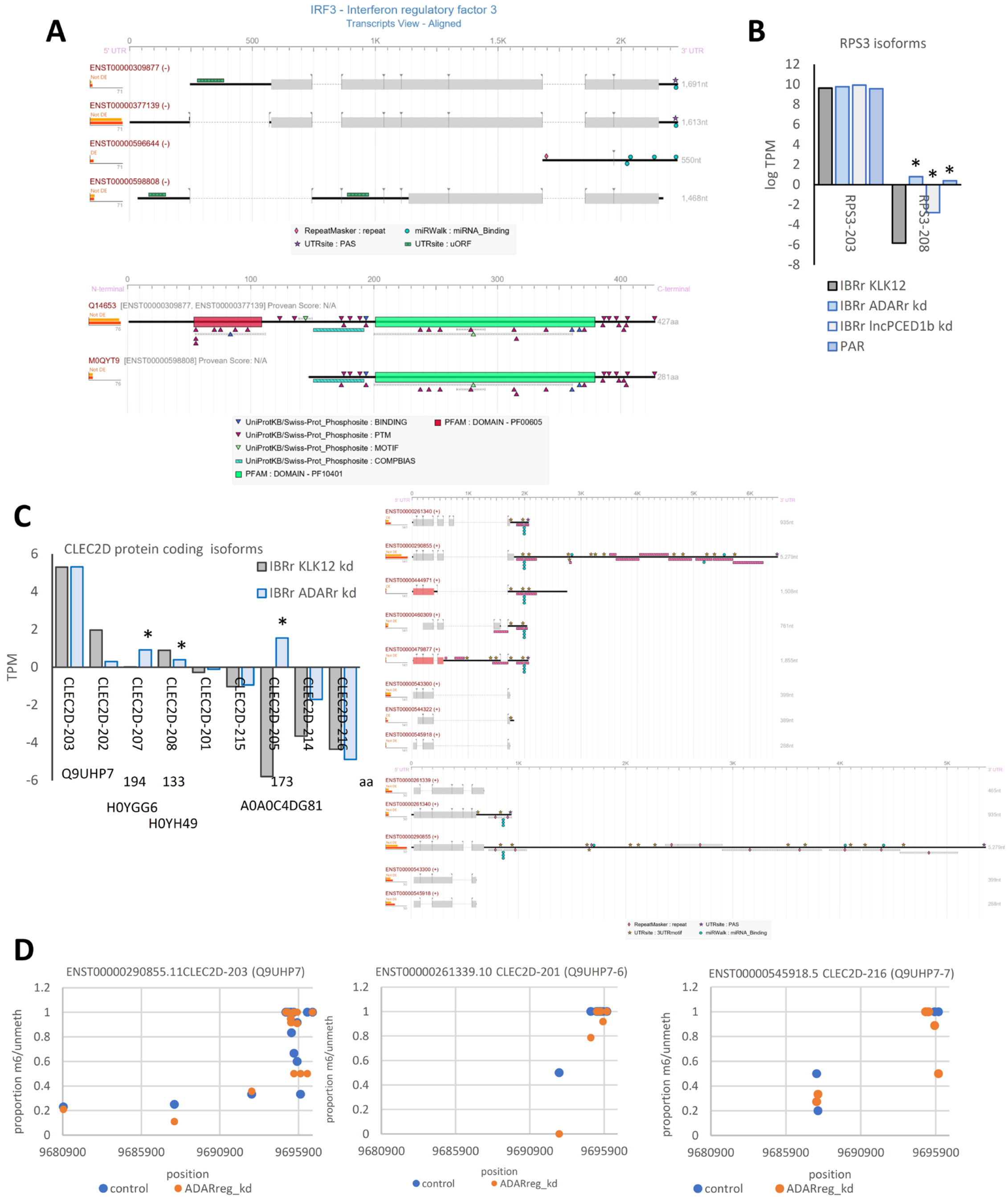


**(A)** TappAS graphical representation of cytosolic IRF3 isoforms with RNA, top, and protein-level annotations, bottom. Colored boxes represent protein domains.

**(B)** RPS3 isoforms measured in total RNAseq performed in parental and ibrutinib resistant VL51 negative control cells and ibrutinib resistant VL51 ADARreg or lncPCED1b kd. * FDR<0.05.

**(C)** Left, CLEC2D isoforms measured in total RNAseq performed in ibrutinib-resistant VL51 negative control cells and ibrutinib-resistant VL51 ADARreg. Labels indicate the predicted protein isoform with corresponding aminoacidic sequence length. * FDR<0.05. Right, TappAS graphical representation of CLEC2D isoforms, as quantified by full-length reads in DRseq, in the nucleus (top) and in the cytosol (bottom).

**(D)** Positional representation of $m^6A$ modulation in the comparison ibrutinib-resistant ADARreg_kd (orange) versus control KLK12kd (blue), along three cytosolic protein-coding CLEC2D isoforms, Q9UHP7 (left), Q9UHP7-6 (middle), Q9UHP7-7 (right).

**Figure S8 (related to Figure 7):** Crosstalk between NK cell and tumor correlated with ADARreg signature in CLL patients.

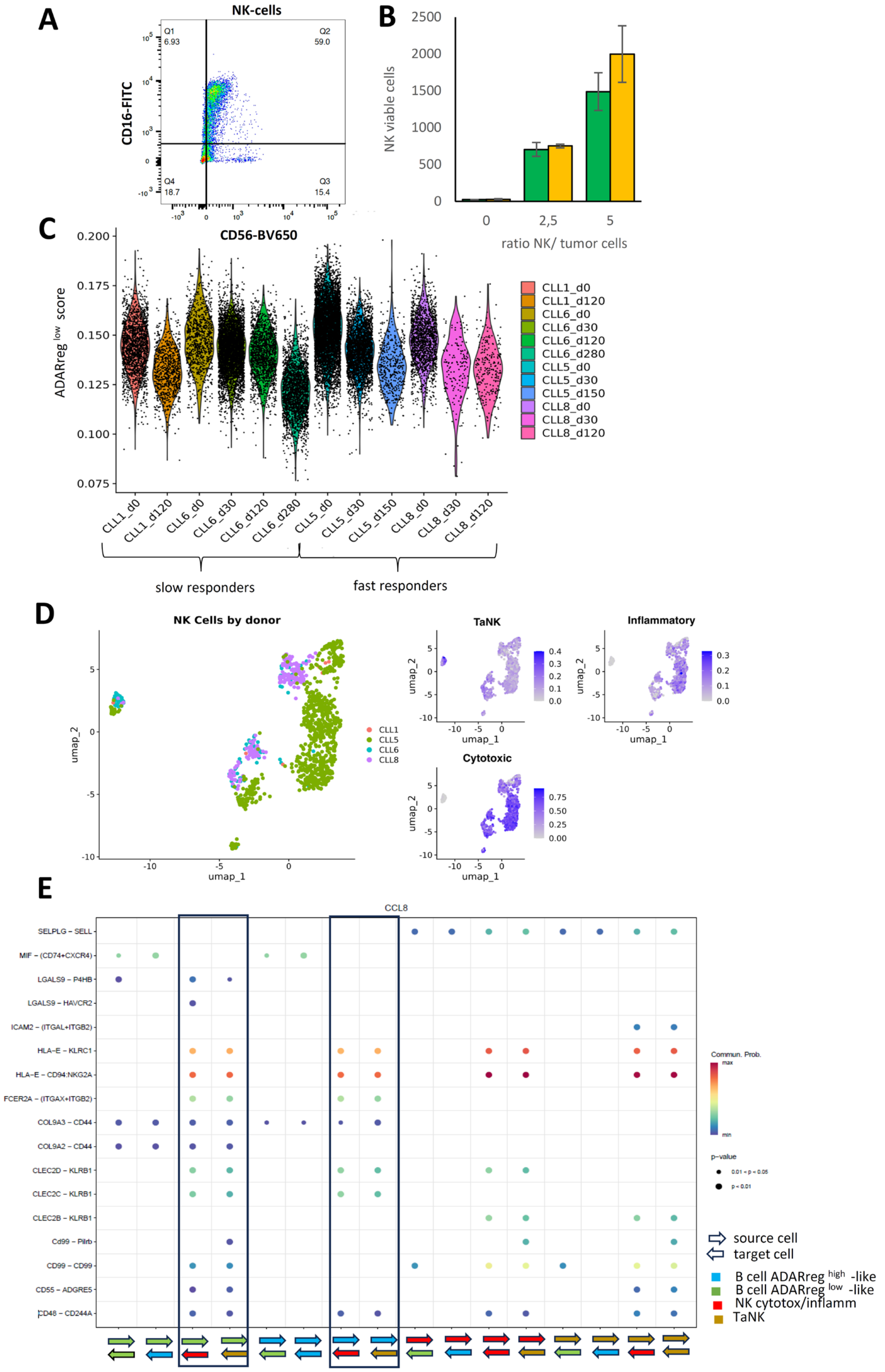

**(A)** Purity check of healthy donor NK cells after magnetic selection by FACS analysis of CD16-CD56 expression.

**(B)** Bar graph representing the absolute number of viable NK cells after 7 days of co-culture with GFP+ mCherry+ ibrutinib resistant VL51, negative control or ADARreg kd. Average of three independent biological replicates. * P <0.05

**(C)** Violin plot representing the distribution over time of B cells in four CLL patients, two fast (CLL5 and CLL8) and two slow ibrutinib-responders (CLL1 and CLL6), based on ADARreg$^{low}$ score.

**(E)** Uniform manifold approximation and projection (UMAP) visualizations of NK cells in the four patients (left) and scored according to tumor-associated, inflammatory, or cytotoxic NK genesets (right).

**(F)** Intercellular communication networks inferred by CellChat analysis between ADARreg$^{high}$-like (blue) or ADARreg$^{low}$-like (green) B cell and cytotoxic or inflammatory NK (red), or tumor-associated NK (TaNK) (brown), in ibrutinib fast responder CLL patient (CLL8).

**Figure S9 (related to Figure 7):** proposed model

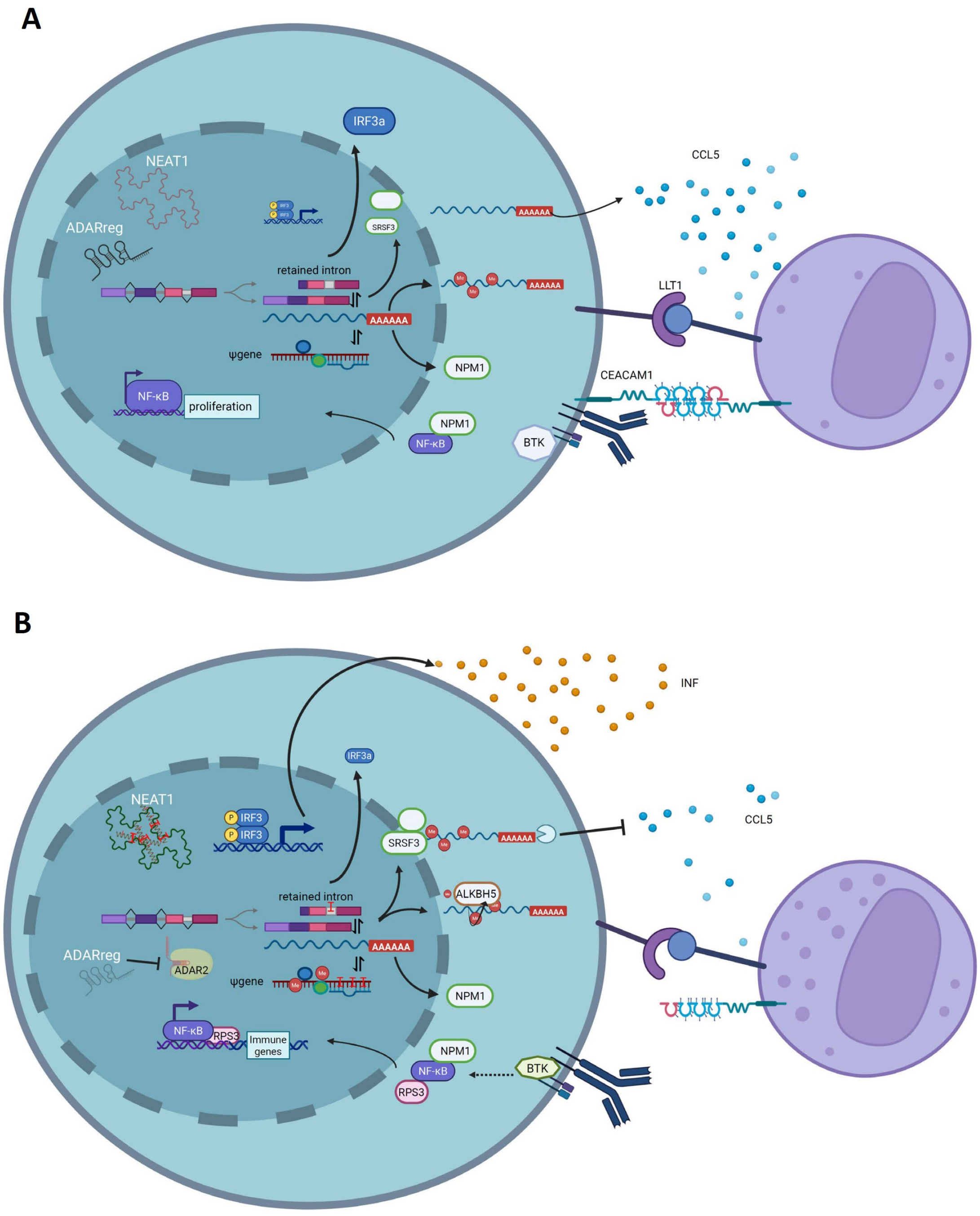


Created in BioRender. Rinaldi, A. (2025) https://BioRender.com/uol128e

**(A)** ADARreg-expressing cells promote an immune-tumor permissive environment through multiple mechanisms, including suppression of IRF3 signaling, production of immunesuppressive chemokines, inhibition of NK-cells activity via LLT1 and CEACAM1 receptors, decoupling of BCR signalling that results in constitutive NF-kB activation.

**(B)** ADARreg silencing increases nuclear ADAR2 activity, leading to RNA editing of intron retain isoforms, that restore IRF3 signalling and induce the expression of RBPs involved in m$^6$A RNA modification (SRSF3 and ALKBH5). These changes, in turn, regulate the expression of chemokines and immune-modulating receptors and the RPS3-mediated-immune signature of NFkB, downstream BCR pathway.

### Table S10 (related to STAR Methods): oligos for CRISPRi sequencing and validation experiments

| | | | | |
|---|---|---|---|---|
| **NGS sequencing primers** | | | | |
| **Name** | **Sequence** | | | |
| P5_VB_ 0 nt stagger | AATGATACGGCGACCACCGAGATCTACACTCTTTCCCTACACGACGCTCTTCCGATCTTCTTGTGGAAAGGACGAAACA | | | |
| P5_VB_ 1 nt stagger | AATGATACGGCGACCACCGAGATCTACACTCTTTCCCTACACGACGCTCTTCCGATCTCTCTTGTGGAAAGGACGAAACA | | | |
| P5_VB_ 2 nt stagger | AATGATACGGCGACCACCGAGATCTACACTCTTTCCCTACACGACGCTCTTCCGATCTGCTCTTGTGGAAAGGACGAAACA | | | |
| P5_VB_ 3 nt stagger | AATGATACGGCGACCACCGAGATCTACACTCTTTCCCTACACGACGCTCTTCCGATCTAGCTCTTGTGGAAAGGACGAAACA | | | |
| P5_VB_ 4 nt stagger | AATGATACGGCGACCACCGAGATCTACACTCTTTCCCTACACGACGCTCTTCCGATCTCAACTCTTGTGGAAAGGACGAAACA | | | |
| P5_VB_ 6 nt stagger | AATGATACGGCGACCACCGAGATCTACACTCTTTCCCTACACGACGCTCTTCCGATCTTGCACCTCTTGTGGAAAGGACGAAACA | | | |
| P5_VB_ 7 nt stagger | AATGATACGGCGACCACCGAGATCTACACTCTTTCCCTACACGACGCTCTTCCGATCTACGCAACTCTTGTGGAAAGGACGAAACA | | | |
| P5_VB_ 8 nt stagger | AATGATACGGCGACCACCGAGATCTACACTCTTTCCCTACACGACGCTCTTCCGATCTGAAGACCCTCTTGTGGAAAGGACGAAACA | | | |
| P7_VB_1-20 | CAAGCAGAAGACGGCATACGAGATNNNNNNNNGTGACTGGAGTTCAGACGTGTGCTCTTCCGATCTCCTTATTTTAACTTGCTATTTCTAGCTCTA | | | |
| | NNNNNNN= sample index | | | |
| **Validated candidates and pgRNAs** | | | | |
| **Alias/Feature** | **Gene Name** | **gRNA_library code** | **sgRNA1 sequence** | **sgRNA2 sequence** |
| EZH2 regulator | LINC00511_A | ENST00000457958.5_gRNA_3030 | GAGAACAAATCAATCCACGGA | ATCAGCTCCGCTTGGAGGCA |
| LOC730338/ADAR regulator | AC011294.3_A | ENST00000451905.3_gRNA_2924 | GACTGGATATGGAATTTCAAG | ACATTTCCTTGTTCCTCTCT |
| lnc-PCED1B-5:28 (alias SLC38A4-AS1) | RP11-96H19.1 | ENST00000611243.1_gRNA_4319 | AAACCACGGGGAATGGAGGT | ACTTGTGGCTGTGGTTATGA |
| negative control | KLK12 | KLK12_gRNA_4607 | GCTAAACGGAGTGGGGAAGG | GGTCTGTCTGGAGCGACAGA |
| **Validated candidates and ASOs** | | | | |
| **Candidate** | **ASO Sequences** | | | |
| AC011294.3-ex1 | CCTTTCATGATATGTT | | | |
| AC011294.3-ex2 | TTCTGCTCTGTACCTA | | | |
| AC011294.3-ex3 | AGACTTCTGGTTGTCC | | | |
| RP11-96H19.1-ex2 | ACCCAGCTGTACTTTG | | | |
| RP11-96H19.1-ex3 | GGGCAGCTGCCTTTAT | | | |
| RP11-96H19.1-ex4 | TCCCTCCCTGGTAAAT | | | |
| NC1-2 | GGCTACTACGCCGTCA | IDT neutral control | | |
| **Primers** | | | | |
| ADARreg ex2-ex3_Fw | ACAGAGACTTGCTTGTTTTGCT | | | |
| ADARreg ex2-ex3_Rev | ACTGTCTTCATCACCTGCGT | | | |
| ADAR reg ex3 Fw | GACAGCAGACCTTCAAGCAC | | | |
| ADAR reg ex3 Rev | GCACGGAACACACTTTCACT | | | |
| LINC00511-202 EX3 Fw | GAGGCCAGTGTTTGTGTGTT | | | |
| LINC00511-202 EX3 rev | ATGGAGAACACGAGCCTTCA | | | |
| lncPCED1b_ex3 Fw | TGCAAGAGTTCCACCCTGAA | | | |
| lncPCED1b ex3 Rev | ACCTTGCCTCCTCTCTTTCA | | | |
| GAPDH ex3 Fw | TCACCAGGGCTGCTTTTAAC | | | |
| GAPDH ex4 Rev | GGGTGGAATCATATTGGAACA | | | |
| KCNQ1OT1 +1150 Fw | AGTCCGCATGGTGGTTTTAG | | | |
| KCNQ1OT1 +1207 Rev | TCAGACTCCCGATCCTCTGT | | | |
| MALAT1_Fw | TGGGGGAGTTTCGTACTGAG | | | |
| MALAT1_Rev | TCTCCAGGACTTGGCAGTCT | | | |

Sequences of pgRNAs cloned in lentiviral vector for stable silencing of elncRNAs, antisense oligonucleotides used for transient silencing, primers to prepare CRISPRi library amplicons for sequencing, and primers to detect transcript expression by qRT.
